# Mapping the convergence of genes for coronary artery disease onto endothelial cell programs

**DOI:** 10.1101/2022.11.01.514606

**Authors:** Gavin R. Schnitzler, Helen Kang, Vivian S. Lee-Kim, X. Rosa Ma, Tony Zeng, Ramcharan S. Angom, Shi Fang, Shamsudheen Karuthedath Vellarikkal, Ronghao Zhou, Katherine Guo, Oscar Sias-Garcia, Alex Bloemendal, Glen Munson, Philine Guckelberger, Tung H. Nguyen, Drew T. Bergman, Nathan Cheng, Brian Cleary, Krishna Aragam, Debabrata Mukhopadhyay, Eric S. Lander, Hilary K. Finucane, Rajat M. Gupta, Jesse M. Engreitz

**Affiliations:** Broad Institute of MIT and Harvard, Cambridge, MA; Department of Genetics, Stanford University School of Medicine, Stanford, CA; BASE Initiative, Betty Irene Moore Children’s Heart Center, Lucile Packard Children’s Hospital, Stanford, CA; Divisions of Genetics and Cardiology, Department of Medicine, Brigham and Women’s Hospital, Boston MA; Department of Biochemistry and Molecular Biology, Mayo Clinic College of Medicine and Science, Jacksonville, FL; Geisel School of Medicine at Dartmouth, Hanover, NH; Faculty of Computing and Data Sciences, Departments of Biology and Biomedical Engineering, Biological Design Center, and Program in Bioinformatics, Boston University, Boston, MA; Cardiovascular Research Center, Massachusetts General Hospital, Boston, MA; Department of Biology, MIT, Cambridge, MA; Department of Systems Biology, Harvard Medical School, Boston, MA; Department of Medicine, Massachusetts General Hospital, Boston, MA; Analytic and Translational Genetics Unit, Massachusetts General Hospital, Boston, MA; Stanley Center for Psychiatric Research, Broad Institute of MIT and Harvard, Cambridge, MA; Currently on leave from the Broad Institute, MIT, and Harvard

## Abstract

Genome-wide association studies (GWAS) have discovered thousands of risk loci for common, complex diseases, each of which could point to genes and gene programs that influence disease. For some diseases, it has been observed that GWAS signals converge on a smaller number of biological programs, and that this convergence can help to identify causal genes^1–6^. However, identifying such convergence remains challenging: each GWAS locus can have many candidate genes, each gene might act in one or more possible programs, and it remains unclear which programs might influence disease risk. Here, we developed a new approach to address this challenge, by creating unbiased maps to link disease variants to genes to programs (V2G2P) in a given cell type. We applied this approach to study the role of endothelial cells in the genetics of coronary artery disease (CAD). To link variants to genes, we constructed enhancer-gene maps using the Activity-by-Contact model^7,8^. To link genes to programs, we applied CRISPRi-Perturb-seq^9–12^ to knock down all expressed genes within ±500 Kb of 306 CAD GWAS signals^13,14^ and identify their effects on gene expression programs using single-cell RNA-sequencing. By combining these variant-to-gene and gene-to-program maps, we find that 43 of 306 CAD GWAS signals converge onto 5 gene programs linked to the cerebral cavernous malformations (CCM) pathway—which is known to coordinate transcriptional responses in endothelial cells^15^, but has not been previously linked to CAD risk. The strongest regulator of these programs is *TLNRD1*, which we show is a new CAD gene and novel regulator of the CCM pathway. *TLNRD1* loss-of-function alters actin organization and barrier function in endothelial cells *in vitro*, and heart development in zebrafish *in vivo*. Together, our study identifies convergence of CAD risk loci into prioritized gene programs in endothelial cells, nominates new genes of potential therapeutic relevance for CAD, and demonstrates a generalizable strategy to connect disease variants to functions.

## Introduction

Genetic variants that influence complex traits are thought to regulate genes that work together in particular biological pathways. Identifying convergence around pathways can help in discovering genes and cellular functions that causally influence disease risk^1–6^. However, it is often challenging to identify such convergence: complex traits often involve contributions from multiple cell types; most risk variants are noncoding and can regulate multiple nearby genes; and it remains unclear which genes work together in which pathways in which cell types^8,16,17^.

GWAS for coronary artery disease have discovered over 300 independent signals^13,14,18^. CAD heritability is significantly enriched in multiple cell types^19,20,21^, including endothelial cells and vascular smooth muscle cells in the vessel wall, and hepatocytes, which influence cholesterol metabolism. At a few individual loci, noncoding risk variants have been shown to regulate the expression of key endothelial cell genes such as endothelial nitric oxide synthase (*NOS3*), endothelin 1 (*EDN1*), and others^20–24^. However, it remains unclear which other genes in CAD GWAS loci might work together in which endothelial cell pathways to modulate disease risk.

To address these challenges, an ideal approach would be to build comprehensive maps of enhancers and gene programs in a given cell type in an unbiased way, such that we could link GWAS variants to the genes they regulate, link genes to the programs they regulate, and determine which of those programs might be relevant to disease risk. Accordingly, we developed a 5-step Variant-to-Gene-to-Program (**V2G2P**) approach (**Fig. 1a**, and **Extended Data Note 1**), in which we:

1. **Identify a cell type and cellular model relevant to disease genetics, through enrichment of disease risk variants in enhancers in that cell type.** Here, we focused on the role of human aortic endothelial cells, by studying telomerase-immortalized (teloHAEC) cells.
2. **Build a map of variant-to-gene (V2G) links in that cell type, to link disease-associated variants to potential target genes.** Here, we consider evidence from variants in endothelial-cell enhancers, as well as coding regions and splice sites.
3. **Build a map of gene-to-program (G2P) links in that cell type, using Perturb-seq**^9–12^ **to systematically knock down candidate disease genes, to identify sets of genes that act together in biological pathways.** Here, we knock down all expressed genes within ±500kb of 306 CAD GWAS signals, and read out the effects of each perturbation with single cell RNA-seq.
4. **Identify “disease-associated programs”, by testing whether the genes with links to risk variants are enriched in (that is, converge on) particular programs.** Here, we find that many CAD GWAS loci converge on the cerebral cavernous malformations (CCM) pathway, whose potential role in coronary artery disease risk has not been characterized.
5. **Nominate “disease-associated genes” as those that are both linked to risk variants and are members of disease-associated programs**. Here, we nominate 41 genes likely to influence CAD risk through effects in endothelial cells, and dissect in detail one new gene, *TLNRD1*, that we show is a novel regulator in the CCM pathway.

**Fig. 1.**
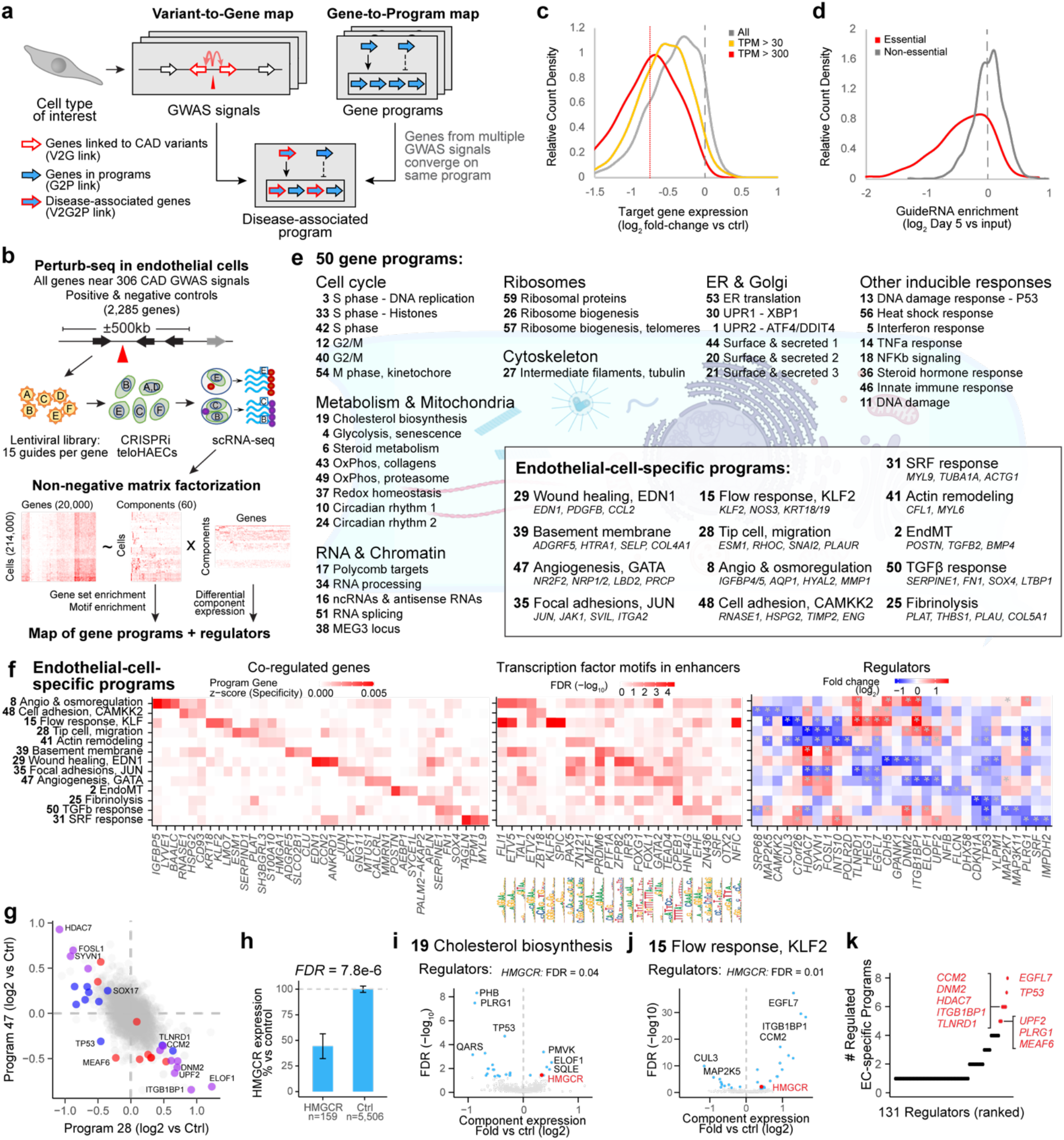
Building a map of gene programs in endothelial cells using Perturb-seq. **a.** Overview of the Variant-to-Gene-to-Program (V2G2P) approach. **b.** Diagram of the approach to create a map of gene programs and regulators using Perturb-seq. **c.** Distribution of knockdown efficiency across target genes (log_2_ expression in cells containing guideRNAs targeting the gene versus in cells containing negative control guideRNAs). Gray line: all targeted genes. Yellow and red lines: Genes expressed at >30 and >300 TPM, respectively. Red dotted vertical line: 40% knockdown (average for 300+ TPM target genes). **d.** Distribution of fitness effects across all guideRNAs (log_2_ ratio of guide frequency in singlet cells from the Perturb-seq experiment after 5 days of CRISPRi induction compared to guide frequency in the original guideRNA library). Guides targeting common essential genes (red) were depleted more frequently than guideRNAs targeting other genes. **e.** 50 gene programs identified from Perturb-seq. Program labels are our annotations based on gene set enrichments and other features of the genes in each program (see also **Supplementary Table 12**). For the 13 endothelial-cell-specific programs, selected co-regulated genes in the program are shown in italics. **f**. Endothelial-Cell-Specific Programs. Heatmaps show the top 3 program co-regulated genes (left, ranked by specificity to the program, see Methods); the top 3 transcription factor motifs in enhancers (middle, ranked by enrichment FDR); and top 3 regulators (right, ranked by fold-change in component expression; see also **Extended Data Fig. 5d**). **g.** Log_2_ fold change in expression of programs 28 versus 47 for each perturbed gene relative to controls. Program 28 (Tip cell, migration) includes co-regulated genes that mark tip cell specification during sprouting angiogenesis (*ESM1, RHOC, PLAUR*), and Program 47 (Angiogenesis, GATA) includes co-regulated genes that are enriched in GATA2 & TAL1 motifs and that include *NRP2*, a co-receptor for VEGF-A, previously shown to act downstream of GATA2^27^). Blue, red, and purple mark genes that are regulators of Program 28, Program 47, or both programs, respectively. **h.** Bar plot showing *HMGCR* gene expression for *HMGCR* guideRNAs versus control guideRNAs. Average knockdown: 53%. Error bar: 95% confidence interval. **i.** Volcano plot shows effects of all perturbations on component expression for Program 19 (cholesterol biosynthesis). *HMGCR* knockdown (red) increased component expression by 26%, FDR = 0.04. **j.** Same as (**i**), for Program 15 (Flow response, KLF2). *HMGCR* knockdown: 33% increase, FDR = 0.01. **k.** 131 perturbed genes that are regulators of at least one endothelial-cell-specific program, ordered by the number of such programs that they regulate. Top 10 regulators are labeled.

In summary, V2G2P defines cell-type specific programs *de novo* using Perturb-seq, combines these programs with enhancer-to-gene maps from the same cell type, and provides an interpretable framework for tracing the path from variant to gene to disease program at individual GWAS loci. This framework builds on recent findings that combining V2G and G2P evidence can identify causal genes with improved precision^3,25,26^.

## A variant-to-gene map in endothelial cells

To implement this V2G2P approach, we collected GWAS signals for coronary artery disease from recent meta-analyses (defining a GWAS signal as a lead variant plus other variants with in linkage disequilibrium (LD), r^2^ > 0.9)^13,14^, and defined a set of “nearby genes” for each GWAS signal to include the 2 closest genes on either side, as well as all genes within 500 kb. We focused on the 228 “non-lipid” GWAS signals that were not associated with lipid levels^13,14^ (see Methods), because lipid-associated signals likely act in hepatocytes or other non-endothelial cell types. Altogether, this yielded 1,942 total candidate genes, with a median of 8 nearby genes per GWAS signal (**Supplementary Table 1**).

We selected telomerase-immortalized primary human aortic endothelial cells (teloHAEC) as a cellular model and collected bulk RNA-seq, ATAC-seq, and H3K27ac ChIP-seq data in resting and several stimulated conditions (+IL1 β, TNFa, VEGFA) to identify expressed genes and candidate enhancers (**Supplementary Table 2**). Variants in teloHAEC enhancers were 11- to-13-fold enriched for CAD heritability by stratified linkage disequilibrium score regression (S-LDSC, **Extended Data Fig. 1a**), supporting the choice of this cellular model.

To link risk variants to genes (V2G), we considered variants for each GWAS signal, and linked variants to genes based on the variant (i) being in a coding sequence or splice site, or (ii) overlapping an enhancer linked to the gene by the Activity-by-Contact model (ABC, which we recently showed performs well at linking noncoding variants to target genes in specific cell types^7,8^), considering the 2 genes per signal with the strongest ABC scores; or (iii) overlapping a chromatin accessible region close to a gene, considering the 2 closest genes to the variant (**Supplementary Tables 1 & 3**). We included enhancers identified in resting and stimulated teloHAEC as well as other previous datasets^8^, to capture many possible endothelial cell states.

We identified 254 of 1,942 nearby genes with a link to a CAD risk variant (“**genes with a V2G link”**), at 125 of 228 non-lipid GWAS signals (range: 1–5 genes per signal). At most GWAS signals, multiple nearby genes had V2G links, because there were multiple candidate variants per signal and/or because individual noncoding variants were linked to more than one gene, consistent with previous observations^8,28^.

## A gene-to-program map in endothelial cells

We next applied Perturb-seq^9–12^ to systematically identify transcriptional programs and their regulators in endothelial cells (**Fig. 1b**). We designed a CRISPRi guide library to inhibit the promoters of all 1,661 expressed nearby genes around lipid and non-lipid CAD GWAS signals, as well as of 624 other control genes, including genes known to regulate endothelial functions and genes not expressed in TeloHAEC, for a total of 2,285 genes (**Supplementary Tables 4 & 5**). We cloned 15 guides per promoter plus 1,000 non-targeting and safe-targeting guides, for a total of 37,637 guides (**Supplementary Table 6**), into a modified CROP-seq vector^12^. We engineered teloHAEC to express KRAB-dCas9 (CRISPR interference (CRISPRi)) under a doxycycline-inducible promoter (**Extended Data Fig. 1b,c**), transduced cells with the guide library at a low multiplicity of infection, harvested cells 5 days after activation of CRISPRi expression, and collected 20 lanes of 10X 3’ single-cell RNA-seq (see Methods).

In total, we obtained data for 214,449 cells expressing a single guide, at an average depth of 10,870 transcriptome-mapped reads with distinct unique molecular identifiers (UMIs) per cell. This dataset included on average 5.7 cells per guide, 85.5 cells per target promoter, and 929,000 total transcript UMIs per targeted promoter (**Extended Data Figs. 2a-f & 3, Supplementary Table 7**). We found that targets were effectively knocked down (**Fig. 1c**), that knockdown of common essential genes decreased fitness (**Fig. 1d**), and that 10.7% of perturbations of expressed targets significantly impacted the transcriptome (**Extended Data Fig. 2g,h,i; Supplementary Tables 8, 9 & 10**).

We applied an unsupervised approach to this Perturb-seq data to discover gene programs (**Fig. 1b**, bottom). First, we used consensus non-negative matrix factorization (cNMF)^29^ to model the gene-by-cell matrix as a linear combination of latent **components** in which each component has a non-negative weight for each gene and has non-negative expression in each cell. We identified 50 components, after optimizing the number of components and excluding components correlated with batch (see Methods, **Extended Data Fig. 4a-d**).

From each component from cNMF, we defined a “**program**”: a set of genes comprised of both “**co-regulated genes**” (the 300 marker genes whose expression is most specific to that component, see Methods) and “**regulators**” (those genes whose perturbations significantly affected the expression of a component compared to negative control guideRNAs, with experiment-wide FDR < 0.05, **Extended Data Fig. 4e**). This analysis defined 50 programs, each including 300 co-regulated genes and from 0 to 35 regulators, and together including 7,564 unique co-regulated genes and 286 unique regulators (for a total of 7,692 “**genes with G2P links**”, **Fig. 1e**, **Extended Data Fig. 5a-c; Supplementary Tables 11**, **12** & **13**).

We annotated the 50 programs through analysis of their regulators and co-regulated genes, and we identified programs representing an array of ubiquitously expressed (“housekeeping”) processes, inducible stress responses, and pathways associated with endothelial cells (**Fig. 1e**). We annotated 13 programs as “endothelial-cell-specific” because they included genes that were on average more highly expressed in endothelial cells than in other cell types (**Extended Data Fig. 5c, Supplementary Table 12**). These endothelial-cellspecific programs included distinct combinations of genes enriched for roles in angiogenesis, extracellular matrix remodeling, barrier function, and the endothelial-to-mesenchymal transition (endoMT), and the promoters of their co-regulated genes were enriched for different transcription factor motifs (**Fig. 1f, Extended Data Fig. 5d,e**). For example, Program 15 (Flow response, KLF2) appeared to correspond to a canonical endothelial cell response to laminar shear stress defined by the known flow-responsive transcription factor *KLF2:* the program was highly enriched for KLF motifs in promoters; included known flow-responsive genes such as *KRT18/19, NOS3*, and *KLF2* itself; and was significantly reduced by perturbations to *MAP2K5* (MEK5), a kinase known to act in the signaling pathway upstream of KLF2 (**Fig. 1f**, **Extended Data Fig. 5f**)^30,31^.

Analysis of program regulators showed expected effects on known pathways (*e.g*., perturbation of HMG-CoA reductase (*HMGCR*) caused upregulation of programs corresponding to cholesterol biosynthesis and KLF2 target genes^32–34^, **Fig. 1h-j, Extended Data Fig. 5f**), and identified cases where programs shared regulators with the same or opposite direction of effect (*e.g*., regulators that increased the expression of Program 28 led to decreased expression of Program 47, **Fig. 1g**). Most regulators significantly affected only a single program, but 31 perturbed genes significantly regulated 5 or more programs (**Extended Data Fig. 5g**), and 10 perturbed genes regulated 5 or more endothelial-cell specific programs, including genes known to have important functions in endothelial cells such as *EGFL7* and *ITGB1BP1/ICAP1* (**Fig. 1k**, see also **Extended Data Fig. 5h**). Taken together, this gene-to-program map represents a wide range of cellular pathways, links upstream regulators to coherent sets of downstream genes, and provides a resource for understanding the functions and potential disease-relevance of genes in endothelial cells.

Using this gene-to-program map from Perturb-seq, we annotated the 228 non-lipid CAD GWAS signals, and found that 883 of 1,942 nearby genes were linked to at least one program, and that all 50 programs included one or more genes near CAD GWAS signals. This is consistent with the notion that, by chance, genes near a GWAS signal will be involved in various biological processes, and this G2P information alone is not sufficient to identify likely disease genes or programs.

## CAD GWAS signals converge on 5 gene programs

Accordingly, we next examined genes with both variant-to-gene and gene-to-program links to look for convergence of CAD GWAS signals onto particular disease-associated programs. To do so, we tested whether the 254 genes linked to non-lipid CAD GWAS signals were enriched among the 300 to 335 genes defining each Perturb-seq program. We conducted separate Fisher’s exact tests for program regulators and co-regulated genes and combined the p-values to create a single score per program (see Methods).

We identified significant enrichment for 5 programs: Program 8 (regulation of angiogenesis & osmotic balance), 39 (basement membrane & platelet recruitment), 35 (focal adhesions, JUN), 47 (angiogenesis, GATA2), and 48 (calcium-dependent cell adhesion) (**Fig. 2a**). Each of these **CAD-associated programs** included 12 to 18 genes with V2G links for CAD, versus 4.5 expected by chance (2.6- to 4-fold enrichment, FDR < 0.05, **Fig. 2a, Supplementary Table 12**). Together, these 5 programs included 41 unique **CAD-associated V2G2P genes** (genes with V2G links and G2P links to CAD-associated programs), including genes near 43 of 228 non-lipid GWAS signals (**Fig. 2b, Supplementary Tables 1 & 15**).

**Fig. 2.**
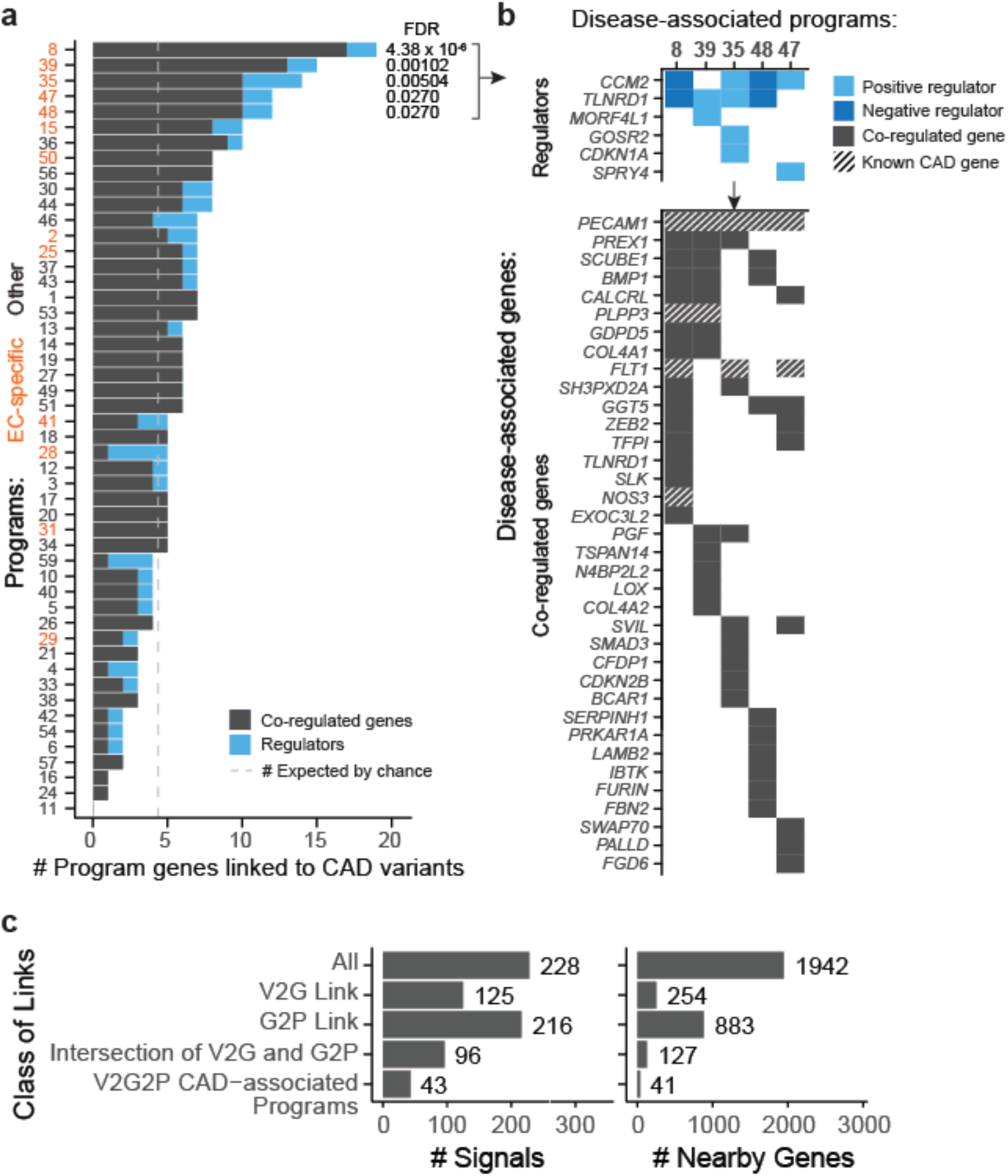
CAD genes converge on 5 programs in endothelial cells. **a**. Identification of CAD-associated programs with V2G2P analysis. The 50 programs are ordered (*y*-axis) by the number of program genes linked to CAD variants (*x*-axis). We define the 5 programs with FDR < 0.05 as CAD-associated programs. Gray dashed line: the number of genes linked to CAD variants that would be expected by chance. Orange labels: endothelial-cell-specific programs. **b.** Relationships among the 41 CAD-associated V2G2P genes and 5 CAD-associated programs. Top: 6 CAD-associated V2G2P genes were regulators of one or more CAD-associated programs (FDR < 0.05). Light blue boxes indicate positive regulators (genes where loss-of-function leads to a decrease in program expression); dark blue indicates negative regulators (genes where loss-of-function leads to an increase in program expression). Bottom: 36 CAD-associated V2G2P genes were co-regulated genes in one or more programs. Gray boxes indicate program membership. Cross hatching indicates genes that were previously known to affect CAD risk through effects in endothelial cells (**Supplementary Table 15**). **c.** V2G2P analysis prioritizes a small subset of genes and GWAS signals compared to either V2G or G2P information alone. Barplots: Counts for signals (left) or nearby genes (right), total (“All”) or those that have: a V2G link, a G2P link, both a V2G link and G2P link to any program, or both a V2G link and a G2P link to a significantly enriched (CAD-associated) program.

Several independent lines of evidence supported the associations of these 5 programs and 41 genes with CAD. (i) All 5 CAD-associated programs included at least 1 of the 8 known genes whose variant-to-gene-to-disease effects in endothelial cells have previously been characterized (**Supplementary Tables 14 & 15**). Program 8 included 4 such genes: *NOS3, PLPP3, FLT1*, and *PECAM1*. (ii) All 5 CAD-associated programs were significantly enriched for CAD heritability by MAGMA^2^, and two were significantly enriched for CAD heritability by S-LDSC^35,36^ (FDR < 0.05, see Methods; **Extended Data Fig. 6a,b**). (iii) The 41 CAD-associated V2G2P genes were highly ranked by an independent gene prioritization method, PoPS^3^, compared to other nearby genes at the same GWAS signals (rank-sum test *P* = 2.5 × 10^−53^, **Extended Data Fig. 6c,d, Supplementary Table 15**). (iv) 9 of the 41 CAD-associated V2G2P genes have previously been found to affect atherosclerosis and/or vascular barrier integrity via studies in mouse models, in a way that is consistent with their acting in endothelial cells (**Supplementary Table 15**).

We made several observations that help to explain the ability of V2G2P analysis to identify disease-associated genes and programs (**Fig. 2c**, **Extended Data Note 2, Extended Data Figs. 6, 7 & 8**). Most notably, at most GWAS signals, neither V2G nor G2P information alone was sufficient to identify likely disease genes: 119 GWAS signals had 2 or more genes with a V2G link (up to 5), and 195 GWAS signals had 2 or more genes with a G2P link (up to 25), including links to all 50 programs. These observations are consistent with the expectation that noncoding variants often regulate multiple nearby genes^8,28^, and that, by chance, a given GWAS signal might have several nearby genes involved in various cellular programs. Combining these two layers of information, however, identified only 5 endothelial cell-specific programs significantly associated with CAD, and, for the 43 signals with V2G2P links to these programs, only 6 had more than 1 linked gene (up to 2). This finding further supports other observations that combining locus-specific variant-to-gene links with genome-wide enrichments for gene pathways can improve the specificity of disease gene identification^3,25,26^.

In summary, our V2G2P analysis supports that CAD GWAS signals converge on specific gene programs associated with disease in endothelial cells.

## The cerebral cavernous malformations pathway regulates CAD-associated programs

Remarkably, all 5 disease-associated programs were regulated in Perturb-seq by the same CAD-associated V2G2P gene—cerebral cavernous malformations 2 (*CCM2*)—and/or by other genes known to act in the same pathway (**Fig. 2b, Fig. 3a,b**).

**Fig. 3.**
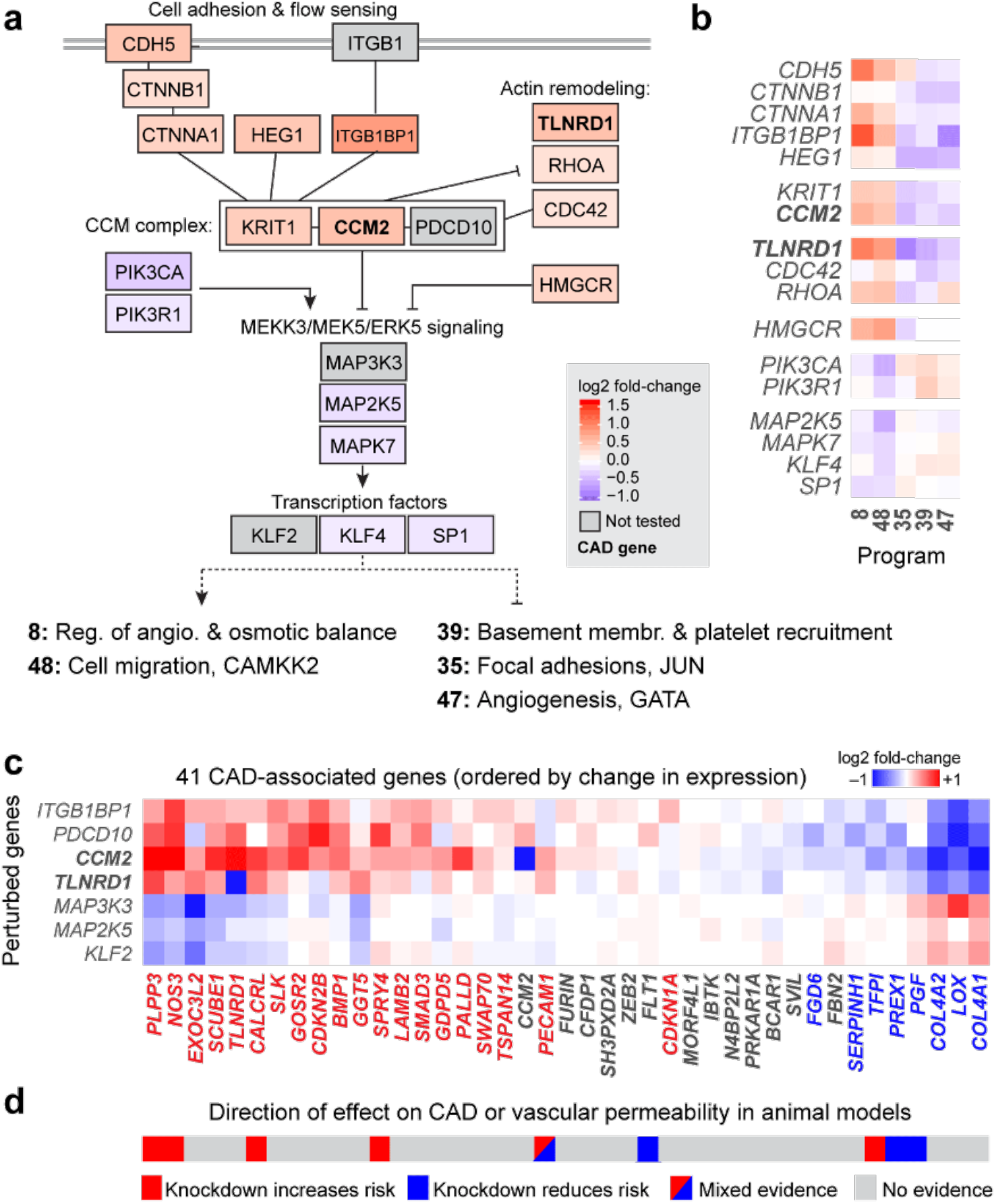
Regulatory connections among CAD genes in the CCM pathway. **a.** The CCM complex and other members of the CCM pathway regulate CAD-associated programs. Color scale: average log_2_ fold-change of effect of the perturbed gene on the 5 programs, with red indicating knockdown leads to increased expression of Programs 8 and 48 and reduced expression of Programs 35, 39, and 47. Solid black lines indicate previously known physical or functional interactions (see Methods). *TLNRD1* is newly linked to the CCM complex via our analysis (see next section). Dotted black lines indicate regulation of the 5 programs. Gray boxes indicate functionally related genes that were not tested in the Perturb-seq experiment. Bold text: CAD-associated V2G2P genes. **b.** Effects of genes in panel A on the 5 CAD-associated programs downstream of the CCM complex. Color scale: log_2_ fold-change on program expression in Perturb-seq. **c.** Effects of perturbing CCM pathway members on expression of the 41 CAD-associated V2G2P genes. Color scale: log_2_ fold-change on gene expression in individual knockdown experiments assayed by bulk RNA-seq (average for two guides to each target). Bold genes: CAD-associated V2G2P genes. Colored text in columns: Genes significantly regulated by one or more CCM pathway perturbation (FDR < 0.05), red: upregulated by upstream signaling gene perturbations or downregulated by downstream gene perturbations, blue: vice versa. **d.** Likely direction of effect of CAD-associated V2G2P genes on atherosclerosis or vascular barrier dysfunction based on prior genetic studies in mouse models (see **Supplementary Table 15** for citations).

The CCM complex—comprised of *KRIT1* (CCM1), *CCM2*, and *PDCD10* (CCM3)—is named as such because loss-of-function mutations in any of these three genes cause cerebral vascular malformations^37–39^, in which inactivation of the CCM complex in cerebral venous endothelial cells activates MEKK3/MEK5/ERK5 signaling and increases downstream activity of KLF2/4^40^. Studies of the CCM complex in both venous and arterial endothelial cells have suggested it acts as a key signaling hub that senses and integrates information about cell-cell contacts, extracellular matrix interactions, and laminar blood flow, and regulates endothelial cell phenotypes *in vitro* such as cell migration and barrier integrity^41^. However, a possible link between the CCM complex and coronary artery disease has not been explored.

In our variant-to-gene analysis, *CCM2* harbored a missense coding variant (rs2107732) that has been associated with a decreased risk of CAD in multiple recent GWASs(odds ratio: 0.92, *P* = 1.53 × 10^−8^)^13,14^. rs2107732 is the lead variant for this GWAS signal (**Extended Data Fig. 9a**), and leads to a valine-to-isoleucine substitution in CCM2 at amino acid 74 — in the PTB domain that is responsible for CCM2 interactions with KRIT1^42^.

In Perturb-seq, knockdown of *CCM2* regulated 4 of the 5 CAD-associated programs at an experiment-wide FDR < 0.05, and the fifth was nominally significant (*P* < 0.05) (–28% to +90% effects on program expression; **Fig. 3b**, **Extended Data Fig. 9b**). Other genes known to interact physically or functionally with CCM2 showed directionally concordant effects on the CAD-associated programs, including *KRIT1*, VE-cadherin (*CDH5*), integrin B1 binding protein 1 (*ITGB1BP1*), alpha catenin (*CTNNA1*), and heart of glass 1 (*HEG1*) (**Fig. 3a,b**, **Extended Data Fig. 9c**). As expected, knockdown of genes known to be repressed by *CCM2* — including MEK5 (*MAP2K5*), ERK5 (*MAPK7*), and *KLF4* — affected the expression of these gene programs in the opposite direction (**Fig. 3a,b**).

Downstream of the CCM pathway, the CAD-associated programs corresponded to distinct sets of genes related to extracellular matrix (ECM) organization, cell migration, and angiogenesis — all processes that have been observed to change upon inactivation of the CCM complex^15,40,41^ and that may have roles in atherosclerosis^43–46^ (**Fig. 3a, b**, **Extended Data Fig. 10**). Program 8 included genes involved in negative regulation of angiogenesis (*IGFBP4* and *IGFBP5*) and osmotic balance (*SLC12A2* and *AQP1*), and included known CAD genes such as *NOS3* and *PLPP3*, which are protective against atherosclerosis^47,48^. Program 48 included genes involved in cell adhesion and migration such as *FSLT1* and *TIMP2*, and was regulated by *MEK5/MAP2K5, ERK5/MAPK7*, and calcium/calmodulin-dependent (*CAMKK2*) signaling. Program 39 expressed genes involved in the basement membrane (*COL4A1/2*) and platelet recruitment (*VWF, SELP*). Program 35 expressed genes involved in focal adhesions (*ITGA2*) and the JAK/STAT signaling pathway. Program 47 also expressed genes involved in angiogenesis including *NR2F2* and *NRP1/2*, including some genes associated with a stalk cell phenotype (*VWF, EHD4*).

To further characterize the co-regulation of the 41 CAD-associated V2G2P genes, we individually knocked down 6 genes in the CCM pathway (*ITGB1BP1, CCM2, PDCD10, MAP3K3, MAP2K5* and *KLF2*) and measured gene expression changes using bulk RNA-seq. 28 of the 41 CAD-associated V2G2P genes were significantly differentially expressed upon CCM pathway perturbation (FDR < 0.05, **Fig. 3c**, **Extended Data Fig. 9d, Supplementary Tables 15, 19**). 8 of these 28 genes have previously been studied in mice to characterize their functions in endothelial cells on atherosclerosis and or vascular permeability (**Supplementary Table 15**), allowing us to assess how changes in gene expression downstream of the CCM complex might relate to disease phenotypes *in vivo*. The direction of effect on disease phenotypes and response to CCM2 knockdown were similar (**Fig. 3d**): of the 5 genes previously shown to maintain vascular barrier function or be protective for atherosclerosis, 4 (*NOS3, PLPP3, CALCRL*, and *SPRY4*) were up-regulated in response to CCM2 knockdown, whereas both of the genes previously shown to promote atherosclerosis or vascular dysfunction (*PGF* and *PREX1*) were down-regulated. One additional gene (*PECAM1*) has been observed to have mixed directions of effect on disease depending on the genetic model (**Fig. 3d**, **Supplementary Table 15**). Thus, down-regulation of the CCM complex leads to changes in gene expression that may be protective for CAD. Interestingly, this is opposite of the direction of effect of the CCM complex on cerebral cavernous malformations, where loss-of-function mutations increase risk^15^.

These findings implicate the CCM complex, together with its associated regulators and downstream transcriptional effects, in controlling endothelial cell functions to influence CAD risk.

## From association to function at *15q25.1*

Among the novel CAD-associated V2G2P genes, we focused further on *TLNRD1* (talin rod domain containing 1), a gene with no known functions in vascular cells, near the 15q25.1 CAD GWAS signal (see also **Extended Data Note 3**, for other novel genes related to CCM signaling). *TLNRD1* was remarkable in that it was the gene that most strongly regulated the CAD-associated programs (**Figs. 2b**, **3b**, **4a, Extended Data Fig. 11a**) and was the gene whose transcriptional phenotype across all 50 programs most strongly correlated with that of *CCM2* (**Fig. 4b**). *TLNRD1* has previously been found to interact with F-actin^49,50^, similar to some other proteins known to interact with the CCM complex (**Fig. 3a**), and to regulate cell migration in a cancer cell line^49^. However, it has not been previously linked to coronary artery disease, the CCM complex, or any function in endothelial cells. Accordingly, we explored whether *TLNRD1* might be a CAD risk gene that acts in the CCM pathway.

**Fig. 4.**
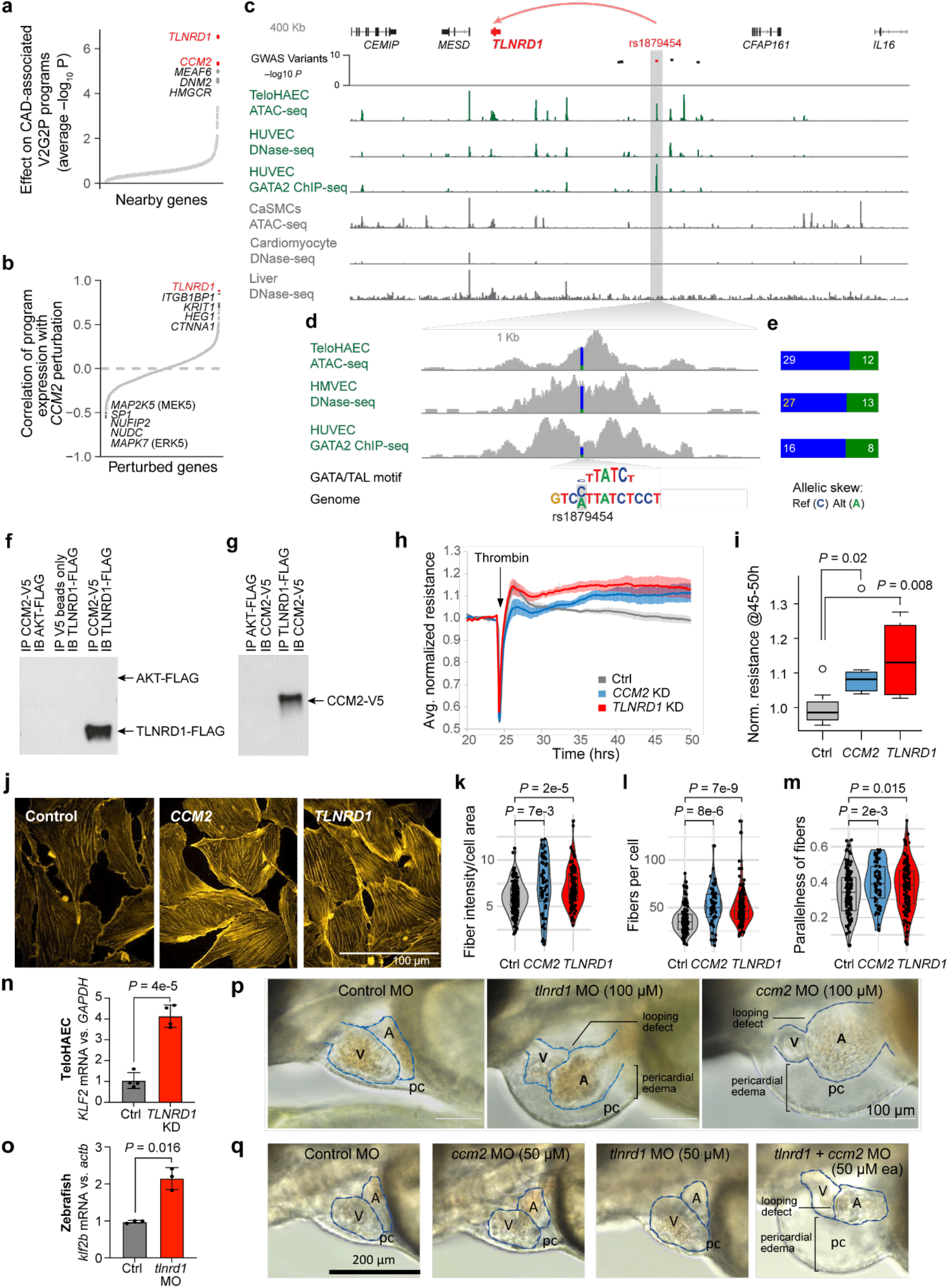
Linking *TLNRD1* to the CCM complex and CAD risk. **a.** 1,503 perturbed nearby genes to CAD GWAS loci, ordered by effect on the 5 CAD-associated programs (average –log_10_ p-value). Labels: top 5 genes. Red: CAD-associated V2G2P genes. **b.** 2,284 perturbed genes ordered by their similarity with *CCM2* perturbation (correlation in log_2_ effects on Program expression). Labels: as in (**a**), for top and bottom 5 perturbed genes. **c.** 15q25.1 CAD risk locus, where rs1879454 is predicted to regulate *TLNRD1* (red arc). GWAS variants: –log_10_ GWAS *P*-value^13^ for variants with LD *R*^2^ > 0.9 with the lead SNP. Green signal tracks: Epigenomic data from endothelial cells. Gray signal tracks: data from other cell types. HUVEC: human umbilical vein endothelial cells. CaSMCs: coronary artery smooth muscle cells. **d.** Zoom-in on the enhancer containing rs1879454. Colored bar in signal tracks indicates read coverage of the reference (C, blue) and alternate (A, green) alleles. Bottom shows the position-weight matrix for a composite GATA/TAL motif and the genome sequence with reference and alternate alleles highlighted in gray. The allele-specificity of chromatin accessibility, measured by DNAse, was also examined in HMVEC (human microvascular endothelial cells, 1.9-fold decrease, *p*-value = 0.0192). **e.** Allele-specific counts for rs1879454 in ATAC-seq in TeloHAEC, DNase-seq in HMVEC, and GATA-2 ChIP-seq in HUVEC. Reads were re-aligned to both reference and alternate alleles to avoid bias toward the reference allele (see Methods). **f.** FLAG-tagged TLNRD1, FLAG-tagged AKT, and/or V5-tagged CCM2 were expressed in HEK293T cells, as indicated. Extracts were co-immunoprecipitated with anti-V5 beads and blotted with anti-FLAG. **g.** As per (**f**), except that extracts were co-immunoprecipitated with anti-FLAG beads and blotted with anti-V5. **h.** Trans-endothelial electrical resistance (TEER), at 4000 Hz frequency, was measured over 50 hrs, for CRISPRi TeloHAEC cells expressing single guides targeting *CCM2* or *TLNRD1*, or non-targeting controls. Thrombin was added 25 hrs after cell seeding, to disrupt cell junctions, and TEER measurements for each well normalized to the average value for the 4 hours before thrombin addition. Data for 2 different guides per target were averaged. N=6 to 8. Ranges: SEM. **i.** Boxplot of normalized TEER signal, from (**h**), averaged for hours 45 to 50 (20-25 hrs post-thrombin). Center line, median; box limits, upper and lower quartiles; whiskers, 1.5x interquartile range; points, outliers. Significance was assessed by two-sided T-test. **j.** CRISPRi TeloHAEC with control, *TLNRD1* or *CCM2* guides were treated with doxycycline for 5 days, then fixed and stained with phalloidin to measure filamentous actin, and 63x confocal images collected (1 well each for 2 separate guides to each target, 25 images per well). Representative maximum-projection images using a fixed intensity range are shown. **k.** Quantitation of actin fiber characteristics in the phalloidin channel from maximum-projection images was performed as per ^54^. For fiber intensity per cell area, integrated intensity in skeletonized fibers was divided by cell area. Boxplots (inset in violin plots), as per (i). N for cells with control, *CCM2* or *TLNRD1* guides was 145, 47, and 117, respectively. **l.** As in (**k**), but showing the number of fibers per cell. **m.** As in (**k**), but showing parallelness of actin fibers. A score of 0 indicates randomly oriented fibers, and a score of 1 indicates all fibers in a cell are parallel. **n.** qRT-PCR for the induction of *KLF2* in TeloHAEC with Cas9-guide nucleofection knock down of *TLNRD1* (or non-targeting guides, “Control”). Signal was normalized to *GAPDH*, and then to the average for controls. **o.** qRT-PCR for the induction of *klf2b* in zebrafish embryos treated with control morpholinos or morpholinos targeted to *tlnrd1*. Signal was normalized to Actin, and then to the average for controls. **p.** In zebrafish embryos, *tlnrd1* knockdown with 100 μM *anti-tlnrd1* morpholino causes cardiac looping defects and severe pericardial edema, similar to *ccm2* knock down, and distinct from 100 μM control morpholino. A: atrium. V: ventricle. pc: pericardial space. **q.** As in (**p**), but showing the synergistic phenotype of 50 μM *tlnrd1* & 50 μM *ccm2* morpholinos, which, individually, showed no phenotype in 41 or 42 zebrafish embryos, respectively, but together showed pericardial edema and looping defects in 103 out of 142 embryos.

The risk variants in the 15q25.1 locus are associated with CAD (lead variant *P*=2.63 × 10^−10^) but not with lipid levels or blood pressure (^13^, **Extended Data Fig. 11b**), and are located in an intergenic region between *CFAP161* (the closest gene) and *TLNRD1* (**Fig. 4c**). Based on our V2G2P analysis, *TLNRD1* was the only gene in the locus with both a variant-to-gene and a gene-to-CAD-associated-program link (**Extended Data Fig. 11c**).

To determine whether a variant in this locus might indeed regulate *TLNRD1* expression in endothelial cells, we examined epigenomic datasets and found that rs1879454 (hg19 chr15:81377717: C (major, risk allele) → A (minor, protective allele); MAF = 0.16) overlapped a chromatin accessible region in teloHAEC and a GATA2 ChIP-seq peak in HUVEC cells. The A allele was predicted to disrupt a GATA motif, and, in cells heterozygous for this variant, the A allele was associated with a 2-fold decrease in allele-specific GATA2 ChIP-seq signal (binomial *P* = 0.0758) and a 2.4-fold decrease in allele-specific ATAC-seq signal (binomial *P* = 0.0058) (**Figs. 4d,e**).

We next explored the functional relationship between TLNRD1 and CCM2. CCM2 is known to physically interact with many other proteins in the CCM complex and pathway^15^. To test if TLNRD1 might physically interact with CCM2, we expressed FLAG-tagged TLNRD1 and V5-tagged CCM2 in HEK293T cells, and found that TLNRD1 immunoprecipitated with CCM2 pulldown, and vice versa (**Figs. 4f,g, Extended Data Fig. 12**).

Knockdown of *CCM2* in endothelial cells *in vitro* has previously been observed to lead to re-organization of the actin cytoskeleton and changes in barrier function^15,41,51^. Both of these phenotypes also occur during endothelial cell responses to laminar shear stress, which is thought to protect against atherosclerosis *in vivo*^44^. We conducted similar tests and found that CRISPRi knockdown of either *TLNRD1* or *CCM2* in teloHAEC led to significant increases in endothelial cell barrier function, as measured by trans-endothelial electrical resistance after disruption of cell-cell contacts with thrombin (**Figs. 4h,i**), and significant increases in the intensity, number and alignment of F-actin filaments (**Fig. 4j-m**).

To determine whether *TLNRD1* has an evolutionarily conserved role in the CCM pathway, we examined *tlnrd1* expression and function in zebrafish. Previous studies have identified a role for zebrafish *ccm2* in heart and vascular development^52,53^. We performed similar experiments and found that knockdown of either *tlnrd1* or *ccm2* with morpholinos led to the same cardiovascular phenotypes that have been previously observed for *ccm2*, including atrial chamber enlargement, cardiac looping defects, and pericardial edema (minimum effective dose: 100 μM; **Fig. 4p, Extended Data Fig. 13b**). Consistent with a function for *tlnrd1* in endothelial cells, *tlnrd1* expression was highest in the brain, notochord, heart, and dorsal vein vasculature (**Extended Data Fig. 13a**). Additionally, *tlnrd1* knockdown led to increased expression of *klf2b* expression, similar to the effect of human *TLNRD1* knockdown on *KLF2* expression in teloHAECs (**Fig. 4n,o**). To further explore whether *tlnrd1* and *ccm2* might function in the same pathway, we tested lower doses of morpholinos against *tlnrd1* and *ccm2*, either alone or in combination. A 50 μM dose of either morpholino alone had no effect on cardiovascular morphology, while 50 μM of both morpholinos led to similar effects as the 100 μM dose of either morpholino alone (73% of embryos, **Fig. 4q**, **Extended Data Fig. 13b**), indicating that knockdown of *tlnrd1* and *ccm2* is synergistic with respect to the cardiovascular phenotypes.

Together, these data indicate that *TLNRD1* is a strong candidate CAD gene, regulates phenotypes relevant to CAD in endothelial cells *in vitro*, and appears to be a previously unrecognized and evolutionarily conserved member of the CCM pathway.

## Discussion

This study demonstrates a strategy to combine variant-to-gene and gene-to-program maps to identify disease-associated genes and programs. In particular, Perturb-seq enabled the creation of an unbiased map of gene programs in endothelial cells that, when combined with variant-to-gene maps, identified convergence of CAD genes onto specific programs related to the CCM complex. This V2G2P analysis identified a subset of CAD GWAS loci likely to act in endothelial cells (at 43 of 306 CAD GWAS signals), nominated causal genes in those loci (41 genes, most of which have not been previously implicated in CAD), and mapped these genes onto the CCM pathway as upstream regulators or putative downstream effectors.

Our results suggest a model for how certain CAD risk variants tune endothelial cell functions to influence risk for coronary artery disease. In particular, variants affecting *CCM2* and *TLNRD1* may down-regulate CCM complex activity, and thereby alter the expression of many other genes linked to CAD risk variants, including up-regulation of the atheroprotective factors *NOS3, PLPP3*^47,48,55,56^, and other genes downstream of KLF2/4, and down-regulation of *PREX1, PGF*, and other candidate disease-promoting genes^57,58^ (**Fig. 3, Extended Data Table 15**). These changes in gene expression may help protect against atherosclerosis by improving barrier function in arterial endothelial cells (**Fig. 4h,i**), and also possibly by acting in a non-cell-autonomous fashion on other cell types in the vessel wall (*e.g*., by production of nitric oxide by *NOS3*). Notably, this proposed role for the CCM complex in CAD (where partial down-regulation appears to be protective) differs from its previously known role in cerebral cavernous malformations (where complete loss-of-function is thought to be pathogenic)^15,40,41^. Further work will be required to understand the physiological functions of the CCM pathway in coronary artery atherogenesis, and to study the functions of the many newly-identified CAD-associated V2G2P genes that may present opportunities for therapeutic interventions.

Certain limitations of this study highlight directions for future study. First, our Perturb-seq experiment identified programs that reflect numerous known functions of endothelial cells (**Fig. 1e**), but certain disease-relevant programs and regulators may have been missed, either because they would only be revealed in certain cell states or conditions (*e.g*., inflammation or hyperlipidemia), because they involve physiological or non-cell autonomous functions that cannot be discovered through Perturb-seq, and/or because their effect sizes were too small to be detected. Studies at greater depth and in additional models *in vitro* or *in vivo* will be required to understand the functions of these genes and programs. In addition, while the V2G2P framework appears to be effective at identifying disease genes and programs, we expect that future studies may be able to improve upon the methods for defining and integrating V2G and G2P links.

In summary, our approach establishes a new path to systematically catalog cellular programs to link risk variants to disease genes. By applying Perturb-seq at scale across many cell types and states relevant to various complex diseases, it should be possible to nominate causal disease genes for a large fraction of GWAS loci and map how they converge on particular cellular pathways. Such a project is becoming increasingly feasible, and would provide a foundation for systematic efforts to leverage human genetic data to discover disease mechanisms.

## Notes

### Competing Interest Statement

J.M.E. is a shareholder of Illumina, Inc. and 10X Genomics. J.M.E. has received materials from 10X Genomics unrelated to this work. All other authors declare no competing interests.

