## Extended Data for "Mapping the convergence of genes for coronary artery disease onto endothelial cell programs"

### Methods

#### Cell culture & creation of CRISPRi TeloHAEC

Telomerase-immortalized human aortic endothelial cells (TeloHAEC) were purchased from ATCC, and grown in Lifeline VEGF endothelial cell media (LL-0005) with 1x Penn/Strep. We chose TeloHAEC because, while immortalized, they maintain important *in vitro* EC functions such as tubing, lipid transport and response to inflammatory stimuli[59,60]. Cells were plated at a density of 0.5-1.0 x $10^6$ cells per 10 cm plate and split before reaching 4 x $10^6$/plate (3 to 4 days). To create the TeloHAEC CRISPRi line, cells were transduced with lentiviral vectors containing 1) dox-inducible (tetracycline operator controlled) dCas9-KRAB-BFP (CRISPRi machinery, which targets epigenetic repressors to efficiently silence enhancers or promoters[61–63], Addgene #85449) and 2) rtTA (tetracycline activator) with a hygromycin marker (Addgene #66810). After hygromycin selection (250 μg/ml for 4 days), cells were treated with 1 μg/ml doxycycline (dox, a stable tetracycline analogue) for 3 days before FACS sorting for the top 15% of BFP positive cells, and after a period in culture without dox, treated again with dox and re-sorted (**Extended Data Fig. 1b**). Diagnostic FACS performed immediately before the Perturb-seq screen showed no leaky BFP expression in the absence of dox, and 93% BFP positive cells in the presence of dox (**Extended Data Fig. 1c**). CRISPRi TeloHAEC were passaged for routine maintenance in the absence of dox. Eahy926 cells (a HUVEC + A549 hybrid line) were purchased from ATCC, and grown in DMEM + 10% FBS. To study responses to CAD-associated cytokines, cells were untreated (control), or treated with 10 ng/ml recombinant human IL-1β (Millipore IL038), 10 ng/ml recombinant TNFα (Millipore GF023), or with normal media lacking VEGF (for TeloHAEC) or supplemented with VEGF (1x concentration from LifeLine VEGF media, for Eahy926), for 24 hours.

#### Bulk RNA-seq

Total RNA was harvested from TeloHAEC (parental or CRISPRi lines) by Qiagen RNeasy kit, DNAse treated (TURBO DNAse 15' 37°C), and purified on MyOne Silane beads. mRNA was purified from 1 μg of total RNA using the NEBNext Poly(A) mRNA Magnetic Isolation module (NEB), processed for RNA-seq library generation using the NEBNext Ultra II RNA Library Kit for Illumina (NEB), and sequenced to a depth of 10 to 30 million reads/library. Reads were mapped to the human hg19 genome build, and counts per gene tables assembled as per [7,8]. Differential expression calls were made using Limma VOOM[64] (for parental TeloHAEC & Eahy926) or edgeR[65] (for single gRNA clones of CRISPRi TeloHAEC). Bulk RNA-seq data is available from the Gene Expression Omnibus (GEO), accession GSE210522.

#### ATAC-seq, H3K27ac ChIP-seq & identification of TeloHAEC enhancers

For ATAC-seq, one well of a 12-well plate (~200,000 cells) was directly lysed using a custom TN5 buffer (33 mM Tris Acetate pH 7.8, 66 mM Potassium Acetate, 10 mM Magnesium Acetate, 16% dimethylformamide & 0.1% NP40). 47.5 μl of lysed cells was added to 2.5 μl Tn5 tagmentation enzyme (Illumina) & incubated at 37°C for 1 hr, and the reaction stopped by addition of 20 μl buffer RLT (Qiagen). Products were purified by addition of 1.8 volumes Ampure XP beads (Beckman-Coulter) & magnetic separation of beads, followed by two 80% ethanol washes, brief drying of pellets & resuspension in 23 μl water. Barcoded ATAC-seq libraries were then generated as described in [7,8], and sequenced to a depth of 10-20 million reads per library. Chromatin immunoprecipitation for histone H3 lysine 27 acetylation (H3K27ac) was performed as described in [7,8]. ChIP-seq libraries were prepared using the KAPA Hyper Prep Kit (KAPA Biosystems). ATAC-seq libraries were prepared in biological triplicate, and ChIP-seq libraries in biological duplicate. For both types of libraries, reads were mapped to the human genome (hg19 build) using Bowtie2, and peaks identified using MACS2, essentially as per [7,8]. Raw and processed data are available on GEO: GSE210489 (ATAC-seq) and GSE210491 (ChIP-seq). Enhancers and their predicted target genes were identified by applying the Activity-by-Contact (ABC) model to these data, using ATAC-seq and H3K27ac ChIP-seq as the measures of enhancer Activity, and using a cross-cell type average of Hi-C maps[7,8] as the measure of 3D enhancer-promoter contact frequency (https://github.com/broadinstitute/ABC-Enhancer-Gene-Prediction [7,8]). We used an ABC fractional score threshold of ≥0.015[8].

**Selection of genes for the Perturb-seq library**

Perturb-seq, which involves knocking down hundreds to thousands of genes in parallel and measuring their effects on gene expression using single-cell RNA-seq, has previously been shown to provide a high-content, unbiased view of cellular programs as represented in gene expression [9–11]. We constructed a library of promoter-targeted CRISPRi guides to all potential causal CAD genes **(Fig. 1b)**. First, we identified all coding genes within a 1 megabase window surrounding the lead SNPs from CAD loci identified in either or both of van der Harst et al.[14] and Aragam et al.[13] that were expressed in TeloHAEC (1+ TPM, from bulk RNA-seq). If fewer than 2 expressed genes were found within 500kb up- or downstream of the lead SNP, the window was expanded to include the closest 2 genes to each side (for a total of 1661 genes). Non-coding genes were generally excluded, unless there was strong evidence for regulatory functions, particularly in ECs. Selected genes with TPM <1 were included, particularly if they were known to be important for CAD in tissues where they were more highly expressed (*e.g. PCSK9*), or were regulated by IL1-beta in bulk RNA-seq data in TeloHAEC (FDR<0.05, fold change >1.3). As negative controls, we included guides targeting 48 coding genes expressed in other cell types but not detectably expressed in ECs, and the 132 expressed coding genes within 1 Mb of 16 randomly-selected lead SNPs associated with Inflammatory bowel disease, Crohn's disease or Ulcerative colitis [66], and which did not overlap with CAD loci. As positive controls, and to aid in connecting candidate CAD genes to known pathways in ECs, we targeted the promoters of an additional 284 genes with known roles in a wide range of CAD-relevant EC functions such as barrier formation, TGF-beta signaling and inflammation, as well as major classes of expressed transcription factors and common essential genes. We also targeted an additional 160 promoters of expressed genes predicted to be regulated by EC enhancers containing fine-mapped variants associated with other disease phenotypes expected to be modulated by ECs (migraine, blood clotting in leg, systolic blood pressure, diastolic blood pressure & mean arterial blood pressure, from UKBB, see **Supplementary Table 16**). This gave a total of 2285 genes, some of which were members of more than one category.

**Guide library production and validation**

sgRNA guides were designed to target promoters of the chosen CAD and control genes (15 guides spanning from -150 to +100 relative to the Transcription Start Site (TSS)), using our established pipeline ( [7,8], https://github.com/EngreitzLab/CRISPRDesigner). We included 400 non-targeting guides (that do not have close matches to any region in the human genome) and 600 safe targeting guides (targeting non-genic regions lacking enhancer marks) [61]. Because TeloHAEC are puromycin resistant, we adapted the CROP-opti vector ([12], Addgene, #106280) for Blasticidin resistance ("CROP-opti Blast"), by digesting the vector with BsiWI and MluI, PCR-amplifying the Blasticidin resistance gene from lenti-dCas-VP64_Blast (Addgene, #61425) with added homology arms, and performing Gibson Assembly (Gibson Master mix, New England Biolabs). To create "CROP-opti-BC-Blast", we added HyPR-Seq barcodes between the WPRE element and the U6 promoter of CROP-opti-Blast, as described in [67]. A pool of oligos encoding the guide sequences, plus extensions with homology to the U6 promoter and downstream

scaffold (TATCTTGTGGAAAGGACGAAACACCG &
GTTTAAGAGCTATGCTGGAAACAGCATAG) was synthesized by Agilent Technologies, and cloned into Crop-Opti-BC-Blast by Gibson assembly and bacterial electroporation as described[61], at an average coverage of 202 transformants per guide. Note that, since the vector was prepared from a single clone, diversity of the HyPR-seq barcodes (which were not required for Perturb-seq) was not preserved. The library was sequenced and shown to include all 37,637 designed guides with relatively equal coverage of each (the difference in count frequency between the top and bottom 10th percentiles of guides was 2.8). A lentiviral library was produced using a standard 3-plasmid protocol[61], at a scale to yield 10ml of virus, stored in aliquots at -80°C, with each aliquot thawed only once.

**Perturb-seq: Experimental procedure**
To transduce this library into CRISPRi TeloHAEC, cells were resuspended in media containing 10 μg/ml polybrene at a density of 1e6 cells per ml, mixed with virus and plated 4ml per well to 6-well plates, centrifuged at 2000 rpm for 2 hrs at 30°C, and incubated at 37°C for 2 hrs before addition of another 4 ml media without polybrene. The next day, cells were harvested and plated to 15 cm plates and treated with 15 μg/ml blasticidin for 4 days. The effective viral titre was determined using this same protocol, and a volume of virus was chosen that gave a final measured 15.7% infection rate (such that most successfully transduced cells have only 1 guideRNA). For the Perturb-seq study, 127.5 million CRISPRi TeloHAEC were transduced and selected for blasticidin resistance, for a coverage of approximately 360 cells per guide (as back-calculated from yield at the first post-blasticidin split, using the 36.7 hr doubling time observed in routine culture) to 461 cells per guide (as estimated from initial number of cells and infection rate). After blasticidin selection, cells were treated with 2 μg/ml dox for 5 days (plating 18e6 cells at each split, to maintain complexity of the library). We reasoned that, since atherosclerotic plaques develop slowly, the longer-term transcriptional effects of causal CAD gene disruption would provide the greatest insights into disease mechanisms. Thus, while we have found that knock down of guide-targeted genes is near maximal after 2 days of doxycycline treatment (inducing the CRISPRi machinery), we treated guide-containing cells with 2 μg/ml doxycycline for 5 days, to measure the longer-term consequences of each perturbation. We also used this same 5-day dox treatment protocol for downstream validation studies (e.g. bulk RNAseq of single guideRNA clones).

The presence of guideRNAs in cells allows multiplets (droplets containing 2 or more cells) to be unambiguously identified, as droplets containing more than one guide. This allowed us to load ~10-fold more cells per 10X Genomics lane than the maximum number recommended in the manufacturer's protocol. Briefly, cells were harvested, resuspended in PBS with 1% BSA, counted, and loaded at 150,000 cells per lane on a 10X Genomics Chromium Controller using a 3' scRNA-seq V3 kit (20 lanes, for a total of 3 million cells). Cells were isolated in two batches, with 6 lanes for the first batch, and 14 lanes, across 2 cassettes, for the 2nd batch, 6 hours later. scRNA-seq libraries were generated using the 10X Genomics protocol, and given lane-specific indexes. From the initial amplified cDNA, we used a two stage PCR protocol to generate "dialout" libraries, for each lane. Because the CROP-seq vector expresses a Pol II polyadenylated transcript that ends just downstream of the guide sequence, the dialout libraries identify the guideRNA sequences associated with each droplet[12]. PCR1 oligos for the guide dialout PCR were: CTACACGACGCTCTTCCGATCT &
GTGACTGGAGTTCAGACGTGTGCTCTTCCGATCTTGTGGAAAGGACGAAACACC, and PCR2 oligos were AATGATACGGCGACCACCGAGATCTACACTCTTTCCCTACACGACGCTC & CAAGCAGAAGACGGCATACGAGAT-8bp index sequence-GTGACTGGAGTTCAG.

**Assignment of guideRNAs to cells**
To get complete information about guide assignments, dialout libraries were sequenced to approximately 40-fold saturation. Guides were identified from read 1 sequences, using Bowtie2 to align dialout reads to a "genome" composed of all 37,637 guide sequences, requiring no-mismatches. Aligning read 1 and read 2 sequences linked gRNA sequences with cell barcodes (CBCs, unique to each bead/droplet) and unique molecular identifiers (UMIs). To avoid low-frequency PCR chimeras, we required that each CBC-UMI-guide combination be duplicated at least 4 times **(Extended Data Fig. 2a)**. We then identified the guides associated with each CBC, and the number of different UMIs for each CBC-guide combination. We selected 4 UMIs for any single guide as the threshold to call a cell as containing a guide **(Extended Data Fig. 2b)**. We defined singlets (one cell & one guide per CBC) as having ≥4 UMIs for the most frequent guide and ≥4x less than this for the 2nd most frequent guide (choosing these thresholds to give a good balance between power to detect transcriptional effects and accuracy in measuring the magnitude of these effects, as described under Selection of Singlet Thresholds, below). Doublets and higher multimers, were cells with ≥4 UMIs for the top guide, and one or more additional guides with more than 1/4 this number of UMIs. The counts of identified singlets, doublets and higher multiplets are shown in **Extended Data Fig. 2c**.

**scRNA-seq data pre-processing and subsetting to singlets**
scRNA-seq libraries were sequenced on two Illumina NovaSeq S4 flowcells, yielding 20,245,734,673 total reads, across all 20 libraries. The FASTQ files were processed on the 10X Cloud to run CellRanger count with the hg38 reference genome. We used the "filtered" features (*i.e.*, cell barcodes corresponding to droplets that contain a cell), and combined the outputs from all twenty 10X lanes into a single genes x cell matrix. This analysis identified 822,156 cell-containing droplets (see **Supplementary Table 7** for other CellRanger output statistics). To measure the effects of individual guides on individual cells, we selected only those CBCs identified in the dialout analysis as corresponding to singlet cells. This identified 214,449 singlets (droplets containing one cell and one guide), defined as 4+ unique molecular identifiers (UMIs) for the top guide and ≥4-fold fewer UMIs for any other guide (**Extended Data Fig. 2c**). This gave an average of 5.7 cells per guide and 85.5 cells per target promoter. Average sequencing depth was 10,870 transcriptome-mapped UMIs per singlet cell, and 929,000 transcript UMIs, across all 15 guides, for each target promoter. Raw and processed data, as well as supplemental files for downstream analyses, are available from GEO: GSE210681.

**Estimation of fitness effects**
To estimate the fitness effects of guides, we compared the relative frequency of all 15 guides to a given target in the original library to the frequency of the same guides in singlet cells, and estimated significance by Benjamini-Hochberg adjusted binomial tests. Essential genes were defined as those that scored as fitness reducing in 5 of 7 tested lines in [68].

**Differential gene expression (DE) analysis & knockdown efficacy**
To measure the differential effects of guides to specific target promoters on individual genes, we used edgeR[65], with settings for scRNA-seq from [69], comparing all singlet cells with guides to each target to all singlet cells with any of the 1,000 non-targeting and safe targeting guides. Expressed genes with fewer than 10 UMI counts across all singlet cells were excluded from the analysis. To control for possible batch effects, we included the 10X lane number as a covariate. For average knockdown efficacy for each perturbation (across all 15 guides), we used the $\log_2$ fold change and p-values reported by edgeR. To measure the knockdown efficacy of individual guides, we performed binomial tests on: the number of transcripts for the guide's target in singlet cells with that guide (hits), all transcripts in singlets with that guide (tests) versus a background frequency of (transcripts to the target in other singlet cells)/(all transcripts in other

singlet cells). Note that with an average of 5.7 cells per guide, assigning significance for knockdown effects of individual guides was only possible for genes with high expression in unperturbed cells (*e.g.*, TPM>100). To identify perturbations with a significant effect on the transcriptome, we used the edgeR results for the 48 negative control promoters (for genes not detectably expressed in TeloHAEC) to estimate the number of DE genes found by chance, at thresholds of nominal *p*-value < 0.01 and fold change > 1.15. Perturbations with a significant effect on the transcriptome (across all 15 guides to each target) were defined as having more DE genes, by these same thresholds, than the 48 non-expressed controls (using binomial tests with a background rate equal to the average DE gene count for controls over all genes tested, and multiple hypothesis correction by the Benjamini Hochberg method).

**Selection of singlet thresholds**

Expression of the CROP-seq guide mRNA in TeloHAEC is lower than in some other cell lines, such as K562 & HEK293T[12] resulting in the absence of a clear gap between noise (low UMI CBC-guide combinations that are likely PCR chimeras) and higher UMI-count true guide reads (**Extended Data Fig. 2b**). We hypothesized that reducing stringency for singlet calls could potentially reduce power to detect perturbation effects on transcription (due to increased noise from mis-calling some true doublets as singlets), or could increase power (by increasing the total number of called singlets analyzed). To test which of these was true, we measured the correlation between differential expression calls for cells with guides to a given target in the full Perturb-seq library versus a smaller pilot library tested in resting TeloHAEC, reasoning that parameters that improved the correlation between these separate studies would also increase the power of the full scale library to detect real transcriptional effects. Information about guides, as well as raw and processed data for this “200 gene” library can be found on GEO, with accession number GSE212396. For the pilot library, we chose singlets with the very stringent threshold of 6 UMIs for the top guide and more than 5-fold less for the next most frequent guide (“6&<5x”). For the full Perturb-seq dataset we chose 4 UMIs for the top guide and equal to or more than 4-fold less than the next most frequent guide (“4&<=4x”, our final applied standard, yielding 214,449 singlets), or the relaxed thresholds “3&<=3x” (284,466 singlets) and “2&<=2x” (389,792 singlets). We identified 37 gene targets that were shared between libraries, and which also showed an FDR<0.1 effect on the transcriptome in the full Perturb-seq 4&<=4x dataset (measured as described above). We then ran EdgeR[65,69] for differential expression testing (cells with guides to each of these 37 targets versus cells with control guides), for each library and singlet definition (pilot 6&<5x, or full library 4&<=4x, 3&<=3x, and 2&<=2x). Then, for all genes called as differentially expressed in either the pilot library or the full library (raw *p*-value < 0.01), we measured the correlation in log2 fold changes between the pilot & full scale data, repeating this analysis for each singlet definition.

Lastly, we measured the difference in correlation coefficients (*R*) between the relaxed threshold comparisons (pilot v. full library 3&<=3x, and pilot v. full library 2&<=2x) and the base comparison (pilot vs. full library 4&<=4x). We found that the median correlation between pilot & full-scale studies significantly improved with the relaxed singlet thresholds (**Extended Data Fig. 14a**, with significance assessed by two-sided *t*-test). This indicates that the increased number of called singlets with the relaxed thresholds increased the power to detect real transcriptional effects, despite an expected increase in doublets mis-assigned as singlets. Plotting change in *R* for each target for the 2&<=2x singlet definition ((*R* for pilot v. full library 2&<=2x) - (*R* for pilot v. full library 4&<=4x), *y*-axis) against the *R* value for the base correlation (between the pilot and the 4&<=4x full library singlet definition, *x*-axis), we found that in all 13 cases where *R* started high (>0.15, likely real correlations between strong transcriptional effects), *R* increased (**Extended Data Fig. 14b**). *R* also increased in all but one case where it started out negative (correcting anti-correlations likely driven by noise). Weak positive base correlations were adjusted up or down, potentially improving true correlations and correcting spurious ones. As such, relaxed singlet thresholds might improve power to detect reproducible transcriptional changes more than is indicated by simple mean differences in *R* values. On the other hand, we found that lower stringencies reduced the apparent knock down effect on these target genes, themselves (**Extended Data Fig. 14c**, median log2 fold changes: -0.53 for 4&<=4x, -0.41 for 3&<=3x and -0.42 for 2&<=2x), likely due to the fact that a mis-called singlet that was actually 2 cells with different guides would show half-magnitude transcriptional effects of each guide. Reduced singlet thresholds also decreased median log2-fold changes for target genes across all targets in the full-scale library (**Extended Data Fig. 14d**, -0.368 for the 4&<=4x singlet definition, and -0.327 for the 2&<=2x singlet definition). Based on these observations, we chose the thresholds of 4 UMIs for the top guide and <=¼ this for the next (4&<=4x), to provide a good balance between overall power and accurate detection of the magnitude of effects.

**Data processing prior to defining gene programs**
To remove noncoding RNA from the analysis, we removed genes with names starting with "LINC" and gene names with patterns starting with two letters and six digits. We retained cells with a minimum of 200 unique detected genes and a minimum of 200 UMIs. We retained genes detected in a minimum of 10 cells.

**Consensus non-negative matrix factorization (cNMF)**
To identify sets of genes that are co-expressed across single cells in a dataset, we used non-negative matrix factorization (NMF). NMF decomposes an input cell x gene UMI count matrix ($X$) into a cell x component matrix ($W$) and a component x gene matrix ($H$), such that $X = W \bullet H + E$, where $E$ is the error term. The cell x component matrix $W$ represents the contribution of each component to the cell's transcriptional profile, and the component x gene matrix $H$ encodes information about gene expression programs. The number of components ($K$) is a hyperparameter defined prior to performing matrix factorization (see below). To account for the fact that the NMF algorithm is a stochastic algorithm that depends on the initial seed, we used the consensus NMF (cNMF) method developed by Kotliar et al[29]. The cNMF method, after normalizing each gene's expression to unit standard deviation, factorizes the normalized matrix multiple times (here, 100 repeats); clusters the components from the repeat runs based on their pairwise Euclidean distances; removes the components that show low similarity to any other component (here, threshold on Euclidean distance = 0.2); defines "consensus components" as the median of each of the component clusters; and recomputes the cell x component matrix $W$ using these consensus components.

**Choosing the number of components for cNMF analysis**
To choose the free parameter $K$ (number of components, **Extended Data Fig. 4**), we defined a set of benchmarking statistics and compared the results of cNMF run for $K$ = [3, 4, 5, 6, 7, 8, 9, 10, 11, 12, 13, 14, 15, 17, 19, 21, 23, 25, 27, 29, 30, 35, 40, 45, 50, 55, 60, 100]. We ultimately chose $K$=60 for all downstream analyses.

We examined the following benchmarking statistics:
(i) Number of unique GO terms enriched in program co-regulated genes (**Extended Data Fig. 4a**, see below)
(ii) Number of unique enriched TF motifs in the promoters or enhancers of program co-regulated genes (**Extended Data Fig. 4b**, see below)
(iii) Number of perturbations significantly regulating any component (**Extended Data Fig. 4c**).
(iv) Error of cNMF (difference between the original normalized data and reconstructed data, calculated by taking the sum of squares of the element-wise difference of the data, **Extended Data Fig. 4d**)

(v) Stability of cNMF (a measure of consistency of the components output from repeated runs, represented by the silhouette score[29], **Extended Data Fig. 4d**)

We chose $K$ = 60 for further analysis, as the number of components that gave a low cNMF error value while near-maximizing each other metric.

**Excluding components associated with batch effects**

We examined whether some components identified by cNMF were likely to represent batch effects. To do so, we calculated the Pearson correlation between each of the 20 batches (*i.e.*, 10X lanes) and the expression of each component across all cells. Based on the distribution of batch x program Pearson correlation (**Supplementary Table 12**), we assigned 10 components with Pearson correlation > 0.15 as likely representing batch effects. We used the remaining 50 components for further analysis.

**Defining co-regulated genes for each program**

We defined 'co-regulated genes' for each cNMF component as the 300 marker genes with the highest *z*-score regression coefficient as defined by cNMF[29]. Essentially, cNMF uses a linear regression model to identify coefficients indicating the number of standard deviations each gene's expression would change with the increased usage of a given component. A component's marker genes, then, are those with the highest "marker gene regression coefficients" (or "specificity scores") for that component, and we selected the top 300 of these marker genes as the set of "**co-regulated genes**" for each gene expression "program" (as defined below).

**Defining regulators for each program**

We tested whether gRNAs targeting a given gene led to a significant change in expression of each component from the cNMF model. We used the Model-based Analysis of Single Cell Transcriptomics package (MAST)[70] to compare the expression of each component in cells carrying gRNAs targeting a given gene vs. cells carrying control gRNAs (1,000 safe-targeting and negative control guides), including 10X lane as a covariate to account for batch effects. We removed the guides present in fewer than 3 singlet cells and the perturbations with fewer than 2 guides. We used the Benjamini-Hochberg method to account for multiple hypothesis testing on the MAST *p*-values (**Extended Data Fig. 4e**), and assigned '**regulators**' of a program as those genes whose perturbation affected component expression with FDR < 0.05 accounting for 140,760 total tests (60 programs x 2,346 perturbations, which includes the 2,285 targeted TSSes, as well as targeted enhancers that were not further analyzed in this study, **Supplementary Table 6**).

To confirm that these FDRs were well-calibrated, we also conducted a simulation-based test. For each perturbed gene, we sampled from the control cells (all singlet cells with non-targeting or safe-targeting guides) the same number of cells, and compared these sampled cells to the rest of the control cells using the same MAST[70] procedure. We identified 0 significant regulators in this approach (**Extended Data Fig. 2i**), indicating that our FDR < 0.05 threshold is a conservative estimate. We also performed the same procedure to estimate the background rate for perturbations called as having a significant effect on the transcriptome using EdgeR[65,71,72]

**Definition and annotation of gene expression programs**

We defined a gene expression **program** as the set of genes comprised of both the 300 "**co-regulated genes**" and the significant "**regulators**" for each cNMF component. We annotated programs based on features of their co-regulated genes and regulators, including: by manual curation of genes with known biological functions, by enrichment of transcription factor (TF) motifs in the promoters and predicted enhancers of co-regulated genes, and by GO term enrichment (see below).

**Identifying motifs enriched in promoters and enhancers**
To identify transcription factors that might regulate program co-regulated genes, we calculated enrichment of human transcription factor motifs in the sequences of the promoter and enhancers of the top 300 genes ranked by component specificity score (**Extended Data Fig. 4c**, **Supplementary Table 17**).

We obtained promoter sequences by taking 500 bp surrounding the TSS as previously annotated[7] and enhancer regions from the Activity by Contact model at an ABC score threshold of 0.015 in the TeloHAEC control condition. For a gene that had multiple enhancers, we counted motif instances across all of its enhancers. To match motifs to sequences, we used HOCOMOCO v11 human full scan motifs (https://hocomoco11.autosome.ru/downloads_v11), and Find Individual Motif Occurrences (FIMO) (https://meme-suite.org/meme/meme_5.3.2/tools/fimo), with the default settings, and *p*-value thresholds of $10^{-6}$ for enhancers or $10^{-4}$ for promoters.

For a given motif and a given program, we counted the number of occurrences of a motif in the promoter sequences of either (i) the top 300 program co-regulated genes, or (ii) all expressed genes in TeloHAEC, and compared these two vectors of motif counts using a *t*-test. We computed enrichment by dividing the program gene's average motif match count by the rest of the expressed gene's average motif match count. We tested all pairs of matched motifs (570 for promoter and 590 for enhancer) x 60 programs, and used the Benjamini-Hochberg method to account for multiple hypothesis testing on the *t*-test p-values.

**Determining enrichment of annotated gene sets in components**
To determine if the gene expression programs align with annotated and publicly available pathways, we tested whether the co-regulated genes in each component were enriched in gene sets from the Molecular Signatures Database (MSigDB). To do so, we used the clusterProfiler R package[73] and MSigDB gene sets[74] (here, the gene sets labeled as "all" for all gene sets and "c5" for GO terms only). We filtered the MSigDB gene sets to only those with more than 3 genes and less than 800 genes. We annotated each program with the gene sets that showed significant enrichment among the program genes (*FDR* < 0.05, **Supplementary Table 18**), and compared the number of gene sets showing significant enrichment as a function of the number of programs *K* **(Extended Data Fig. 4a)**.

**Defining endothelial-cell-specific programs**
To annotate programs as "endothelial-cell-specific", we analyzed the degree to which program co-regulated genes were expressed in endothelial cells versus other cell types. We took gene expression transcript per million (TPM) data across all available cell types from FANTOM5 and calculated the expression z-score of each gene across all cell types. To give each gene an endothelial-cell specificity score, we calculated the average of all z-scores for a gene across endothelial cell samples. We defined endothelial-cell specificity scores for each program as the average of the 300 co-regulated genes' specificity scores, and selected 0.19 (90% percentile) as the threshold to call programs as "endothelial-cell-specific" (**Extended Data Fig. 4c, Supplementary Table 12**).

**Variance explained by all cNMF components**
To quantify the fraction of variance explained by all 60 programs jointly, we compared the residual variance in the dataset after subtracting the consensus matrix factorization to the total variance in the dataset:

$$V = 1 - \frac{Var(X - WH)}{Var(X)},$$

where $X$ is the (cell x gene) normalized data matrix input to cNMF, $W$ is the (cell x program) usage matrix, $H$ is the (program x gene) spectra or weight matrix, and matrix variance is defined by summing the column- or gene-level variances:

$$Var(X) = \sum_i Var(X_j).$$

Note that cNMF normalizes the input data so each $Var(X_j) = 1$.

**Variance explained by individual gene programs**
To rank gene programs by variance explained, we devised a method to quantify variance explained by NMF or cNMF components separately. For the $k$'th program $H_k$, we consider the effective matrix decomposition given only this program; the effective usage matrix $B_k$ in this case is given simply by orthogonal projection or ordinary least squares: $B_k = XH'_k/||H_k||^2$, where the prime indicates transposition. We then define the variance explained in terms of the residual fraction as above:

$$V_k = 1 - \frac{Var(X - B_k H_k)}{Var(X)}.$$

Our method may be generalized to any set of programs, but with more than one program the effective usage matrix must be obtained by nonnegative least squares (a single iteration of NMF).

**Defining variants in CAD GWAS signals for variant-to-gene analysis**
CAD lead GWAS variants were derived from both Aragam et al.[13] and Harst et al.[14]. We excluded lead variants from Harst et al. if the variants were in strong LD ($r^2 \geq 0.7$) with an Aragam et al.[13] lead variant or were ≤5Kb away from an Aragam et al. lead variant. An LD-expansion was performed to include variants that are both within a 1 Mb window of, and are in strong LD ($r^2 \geq 0.9$) with the any of these lead GWAS variants in 1000 Genome European ancestry (plink --ld-window-kb 1000 --ld-window 99999 --ld-window-r2 0.9). For each lead variant, we also included variants prioritized through functionally informed fine-mapping (PIP ≥ 0.1) in either study[13,14]. We defined a "**GWAS Signal"** as this collection of variants around, and including, each lead variant.

**Identifying CAD variants associated with lipid levels**
We classified CAD GWAS signals as "lipid" or "non-lipid" based on their association with lipid levels in other GWAS studies, because the CAD GWAS signals also associated with lipids are presumed to act through non-endothelial cells such as hepatocytes. For lead signals included in Aragam *et al.*[13], we defined a CAD GWAS signal to be associated with lipids if the lead variant was linked to "LDL-direct", "Triglycerides", "Cholesterol", "HDL-cholesterol", "Apolipoprotein A", "Apoplipoprotein B", "HDLC" or "LDLC" in the phenome-wide association scan (PheWas) conducted by Aragam et al.[13] For GWAS signals exclusively nominated by Harst et al[14], we used a different procedure in which we considered a signal to be associated with lipids if its lead variant was associated ($P < 5 \times 10^{-8}$) with HDLC, LDLC, TG, ApoA, or ApoB based on GWAS from the UK Biobank (Hilary Finucane and Jacob Ulirsch: https://www.finucanelab.org/data). We refer to the remaining GWAS signals not associated with lipid levels as "non-lipid CAD GWAS

signals”, and focused on this subset of signals as cases where CAD variants might plausibly act in endothelial cells.

**Linking variants to genes**

We used a combination of variant-to-gene methods to identify a list of genes linked to CAD variants that could plausibly act in endothelial cells. At each CAD GWAS signal, we considered as candidate genes at least two genes upstream or downstream of the lead GWAS SNP, and all the genes within +/- 500Kb of the lead variant to be potentially regulated by the GWAS signal. We focused our analysis on protein-coding genes and excluded long noncoding RNAs (“^LINC”), gene isoforms (“-AS”), microRNAs (“^MIR”), small nuclear RNAs (“RNU”), and genes of uncertain functions ("^LOC"). To link CAD variants to genes, we intersected the variants with ABC enhancers[8] in endothelial cells to identify the top two genes most likely to be regulated by each variant (highest 2 ABC fractional scores over 0.015). Specifically, we used ABC data, for enhancers and predicted target genes, from TeloHAEC and Eahy926 (control, or treated with IL1$\beta$, TNF$\alpha$ or VEGF, this study), and from prior ABC analysis of HUVEC (‘endothelial_cell_of_umbilical_vein_Roadmap’, ‘endothelial_cell_of_umbilical_vein_VEGF_stim_12_hours-Zhang2013’, and ‘endothelial_cell_of_umbilical_vein_VEGF_stim_4_hours-Zhang2013’ datasets from [8]). To account for cell state-specific regulation that was not predicted by ABC, we also intersected candidate CAD variants at each signal with ATAC peaks and considered the 2 genes closest to variant-containing peaks as plausibly regulated. We also linked variants to genes if the variant was in a coding sequence or within 10 bp of a splice site annotated in the RefGene database (downloaded from UCSC Genome Browser on 24 June 2017)[75]. We confirmed that these candidate CAD variants were significantly enriched for matching any or all of these criteria (**Extended Data Fig. 6e**). We identified 254 candidate CAD genes, defined as “**genes with V2G (variant-to-gene) links”**, at 125 of 228 non-lipid CAD GWAS signals (**Supplementary Table 1**).

**Identifying CAD-associated programs via variant-to-gene-to-program analysis**

We developed an approach to identify gene programs likely to affect CAD risk through functions in endothelial cells. To do so, we tested whether the 254 genes with V2G links (between CAD variants and enhancers/coding regions in endothelial cells) were enriched in each Perturb-seq program. Specifically, we performed a one-tailed Fisher exact test separately for co-regulated genes and for regulators. For co-regulated genes, we constructed a contingency table for whether a gene is a co-regulated gene (out of 17,472 expressed genes) and whether a gene has a V2G link. For regulators, we constructed a contingency table for whether a gene is a regulator (out of all perturbed genes) and whether a gene has a V2G link. We then multiplied the p-values from co-regulated gene and regulator Fisher exact tests together to get a final program enrichment p-value. We use Benjamini-Hochberg method for multiple hypothesis correction across all 50 non-batch programs. 5 programs showed significant enrichment by this method (FDR < 0.05: Programs 8, 35, 39, 47, 48), referred to as “**CAD-associated programs**”.

**Defining CAD-associated V2G2P genes**

We defined “**CAD-associated V2G2P genes**” as those 41 genes that were both (i) a gene with a V2G link to a CAD variant and (ii) a member of one of the 5 CAD-associated programs (as a regulator and/or co-expressed gene). The 41 genes were linked to 43 GWAS signals due to cases where independent GWAS signals are linked to the same gene.

**Identifying enriched programs via MAGMA**

We tested whether the co-regulated genes in each program were significantly enriched near variants associated with CAD using MAGMA[2]. To do so, we took the CAD summary statistics from Aragam et al.[13] (https://data.mendeley.com/public-files/datasets/2zdd47c94h/files/5b4eb0d7-96e8-4c7e-b109-046107ebd480/file_downloaded), and used the MAGMA --annotate function to summarize CAD association p-values for variants within a 50kb window of all human genes, using the 1000 genomes European reference data for base allele frequencies (https://ctg.cncr.nl/software/MAGMA/ref_data/g1000_eur.zip). We then ran MAGMA to test for enrichment of CAD heritability within 50kb of the top 300 program genes, and corrected for multiple testing (60 components) using the Benjamini-Hochberg method.

**Identifying programs and cell types enriched for CAD heritability via stratified LD score regression**

We used S-LDSC to estimate the enrichment of CAD heritability linked to program genes and to enhancers in TeloHAEC. While the original implementations of S-LDSC linked variants to genes based on genomic distance[35,76], we additionally required that variants either overlap exonic regions of the gene or overlap nearby candidate enhancers in endothelial cells (as in [26,36]). In particular, for co-regulated genes in each program, we derived an annotation for S-LDSC by including exonic regions (exons from transcripts with Ensembl_canonical, appris_principal, appris_candidate, or appris_candidate_longest tags, as indicated in the GENCODE v38lift37 annotations) as well as endothelial *cis*-regulatory elements derived from snATAC-seq[77], from which we merged the 9 adult and 8 fetal sets of endothelial peaks into a single annotation, and for each geneset included all peaks within 50kb of the gene starts and ends. For all peaks, we first converted coordinates from the GRCh38 to the GRCh37 reference assembly using UCSC LiftOver, discarding peaks that could not be converted. To estimate the enrichment of CAD heritability in TeloHAEC enhancers, we required the variants to overlap enhancers predicted by ABC from ATAC-seq and H3K27ac ChIP-seq data in TeloHAEC under control conditions or treated with IL1$\beta$, TNF$\alpha$ or VEGF (ABC score > 0.015). For each set of variants (programs or TeloHAEC enhancers) we ran S-LDSC using 1000G EUR Phase3 genotype data to estimate LD scores, baseline v2.2 annotations as recommended by the LDSC developers[78], and HapMap 3 SNPs excluding the MHC region as regression SNPs. We ranked programs by their enrichments and reported the *p*-values of these enrichments (**Extended Data Fig. 6b**).

**Polygenic Priority Score (PoPS)**

PoPS is a method to nominate likely causal genes in a GWAS locus, which prioritizes genes based on their being members of many gene sets enriched for heritability genome-wide[3]. We applied PoPS to summary statistics from Aragam *et al.*[13] using the predefined set of gene sets as previously described[3] (**Extended Data Fig. 6c,d**). For each GWAS signal, we calculated the PoPS rank among "nearby genes" (2 to either side of the lead SNP, and all within +/-500kb). Previously we have shown that genes with the highest PoP score in the locus are strongly enriched for likely causal genes, as identified by fine-mapped coding variants[3], and that this enrichment increases when further focusing on genes that are both the closest gene and have the highest PoP score. In this analysis, we did not use any features from Perturb-seq and, as such, this method represents an entirely independent method that validates the high likelihood of causality of the set of CAD-associated V2G2P genes.

**Defining gene expression programs for cells carrying control guides**

To examine the gene programs in normal, unperturbed teloHAECs, we used the same analysis pipeline on the subset of cells carrying control guides (5,506 cells). We used cNMF to discover *K*=60 components, and defined 60 "control programs" based solely on the 300 co-regulated genes defining each component (because control guides did not target any genes, so there was no regulator information). Of the 60 programs, 4 programs correlated with batch (Programs 2, 17, 22, 41). We compared the program co-regulated genes between control cells and full library programs (**Extended Data Fig. 7d**). Control program 10 highly overlapped with full library programs 8 and 39. The four control programs that correlated with batch also had high overlap in co-regulated genes with the full library's batch programs. We then utilized the V2G2P approach to prioritize these programs, and found that none of the control programs was enriched for genes with V2G links (**Extended Data Fig. 7e**).

**Identifying genes in the CCM pathway**

**Fig. 3** shows a curated set of genes previously reported to interact physically or functionally with the CCM complex and/or downstream ERK5/MEK5 signaling[41,51,53,79–82], plus one additional gene (*TLNRD1*) that we identify here as a member of the CCM pathway. These genes were manually selected through an iterative process involving examining genes known to interact with the CCM complex and that were found to regulate the enriched programs in Perturb-seq.

**Allelic imbalance analysis for a variant linked to *TLNRD1***

We calculated allelic imbalance in ATAC-seq and ChIP-seq signal for the rs1879454 variant, accounting for any mapping bias toward the reference allele following methods previously described[83]. Specifically, we created two reference genome FASTA files that harbored the reference or alternate alleles at rs1879454; aligned ATAC-seq data to both genome files; selected reads that overlapped the variant coordinate; and used PySuspenders[83] and PySAM (https://github.com/pysam-developers/pysam) to assign and count reads that uniquely aligned to one or the other allele. We applied this procedure to ATAC-seq data from TeloHAEC and the ENCODE datasets ENCSR000EVW (GATA2 ChIP-seq on HUVEC) and ENCSR000EOB (DNase-seq and DGF on HMVEC-dLy-Neo).

**Generation of single-guide CRISPRi TeloHAEC derivatives**

Paired oligos for individual guides (newly-designed, as described for the Perturb-seq library, or with the best KD efficacy in Perturb-seq) were annealed and cloned into the BsmBI site of a CROP-Opti-Blast vector (plasmid available upon request), which were then used to generate lentivirus (as per [61]). CRISPRi TeloHAEC were infected with each virus, in separate wells, and selected for blasticidin (15 μg/ml 4 days), before 5 day dox induction and analysis by bulk RNA-seq, fluorescence imaging or physiological assays. Guides (TargetGene_CloneIndex: ForwardSequence) were: CCM2_C2: GGCAAGAAGGTGAGCGTGCG, CCM2_F6: GAGCCGCTACATGCTCGACCC, CDH5_B8: GCCAGCTGGAAAACCTGAAG, CDH5_D5: GTTGGACTGCCTGTCCGTCCA, ITGB1BP1_C7: GAAGGCCGCGGCACTCCCACG, ITGB1BP1_G8: GAAGTCCGCAACCCGGGGAT, KLF2_C9: GGACCCGGGGAGAAAGGACG, KLF2_G10: GCCGCGGTATATAAGCCGGC, MAP2K5_A11: GCCGAGGCCGCGCGGACTGG, MAP2K5_B5: GTCTGCCCCACCCGGAGACAC, MAP3K3_A4: GTTCCTGAGGTGGAGAACGG, MAP3K3_C3: GCCAATAACAAGAAGGAAGT, MEF2A_C10: GCGGCGCGAAGCGCTGGTGG, MEF2A_H10: GACTGAATTATCCTCTCGGT, Negative_control_B6: GCAACGGTGTACCGCGGATC, Negative_control_D2: GTGGTTCACAACCGGACCCA, Negative_control_D8: GGTGGTTCGGTTTGCGTGGCC, Negative_control_F4: GCTGGGCGGACGTTGGGATA, NFAT5_D4: GGCCTCGCTTCCTGCCGGCG, NFAT5_D7: GGTCCCCGTCCCGCCGGGGG, PDCD10_D11: GACCGAGCAGAAGAGGTCTA, PDCD10_G1: GCCGCTTTACGCCACTCGCGT, TLNRD1_B3: GTGGCTGCGCCGCCGCCCGCA, TLNRD1_D12: GCCTCCGGCAGCCCCTGCGGG.

**Ribonucleoprotein-based CRISPR/Cas9 genome editing**

For some experiments, we used Synthego's ribonucleoprotein (RNP) technology as an orthologous method to knock down target genes, as previously described [84]. Briefly, TeloHAEC were nucleofected with Synthego's Gene Knockout Kit v2 for non-targeting negative control, *CCM2,* or *TLNRD1* using the Lonza 4D-Nucleofector system. For each nucleofection reaction, we used 150,000 cells with 20 pmol of Cas9 and 50 pmol of sgRNA. The cells were then nucleofected (program CA-210) using SG cell line nucleofection solution (Lonza; V4XC-3024). The nucleofected cells were seeded in TeloHAEC culture medium, and harvested 48 hrs later for RNA extraction for qRT-PCR analysis to measure gene knockdown efficiency and perturbation effects.

**Co-immunoprecipitation of CCM2 and TLNRD1**
HEK293 cells were transfected with V5-tagged CCM2, Flag-tagged TLNRD1 and/or Flag-tagged Akt1, as indicated in **Figs. 4f,g**, using FuGENE (E2311, Promega). Two days after the transfection, cell lysates were extracted with IP lysis buffer (87787, Thermo Scientific) and immediately used for co-immunoprecipitation. Immunoprecipitation was carried out using magnetic beads (88805, Thermo Scientific) conjugated with 5 μg of either anti-V5 (13202, Cell Signaling Technology) or anti-Flag (F1804, Millipore Sigma) antibody. Cell lysates were incubated with the antibody-conjugated beads for 20 mins at room temperature. Beads were then washed three times with IP lysis buffer, and precipitants were eluted using 2xLDS sample buffer (NP0007, Thermo Fischer Scientific). Precipitants and total lysates were immunoblotted with anti-V5 (13202, Cell Signaling Technology, ab27671, Abcam), then stripped (21059, Thermo Scientific) and re-probed with anti-Flag (F1804, Millipore Sigma), or vice versa. The Akt1-FLAG vector is Addgene #9021. CCM2-V5 (ccsbBroad304_04281) and TLNRD1-V5 (ccsbBroad304_03872) vectors were obtained from the Broad Institute Gene Perturbation Platform[85]. For TLNRD1-FLAG, cDNA sequences were amplified from the TLNRD1-V5 vector using primers that incorporated an in-frame FLAG tag, and cloned into the pcDNA3.1 backbone.

**Trans-endothelial electrical resistance (TEER) measurements**
For TEER measurements, we used the ECIS Z-Theta instrument from Applied BioPhysics in the 96-well plate system (Applied BioPhysics; 96W10idf). CRISPRi TeloHAEC expressing individual guides to *TLNRD1, CCM2*, or non-targeting guides (2 guides each) were treated for 5 days with 2 μg/ml doxycycline. A gold electrode-containing 96-well ECIS plate was incubated at 37°C and 5% $CO_2$ with culture media for 30 min to equilibrate before coating with 2.5 mg/mL fibronectin in 0.1 M bicarbonate buffer at pH 8.0. Then, the coated wells were inoculated with 45,000 cells in 100 mL media. An additional 100 mL of media was added to each well before initiating the measurements at 4000-Hz AC. At 25 hours, after the cells formed a confluent layer, the culture media was replaced with 200 mL of fresh culture media with 1 U/mL thrombin to disrupt cell-cell junctions, and measurements continued until 50 hrs to observe cell junction recovery after thrombin treatment.

**Fluorescence imaging and quantitation of TeloHAEC**
For quantitation of actin fiber characteristics, CRISPRi teloHAEC expressing individual guideRNAs (targeting *CCM2*, *TLNRD1*, or negative control) were treated with 2 μg/ml doxycycline for 5 days. Cells were fixed *in situ* with by addition of paraformaldehyde to 3.2% for 30 mins at 37°C, washed with PBS, permeabilized by addition of PBS with 0.1% triton X100 for 15 mins at room temperature, washed with PBS and stained with PerkinElmer Cell Painting dyes (Phenovue Fluor 568 - Phalloidin, Phenovue Fluor 488 - Concanavalin A, Phenovue Hoechst 33342 Nuclear Stain & Phenovue 512 Nucleic Acid Stain) according to the manufacturer's instructions. Cells were imaged in four channels as described in [86] on a Perkin Elmer Opera Phenix Imaging System-106513, confocal 63x magnification with 1x binning. The stacks of images for the Phalloidin and Hoechst channels were converted to single images using maximum projection, output ranges standardized, and images exported. Cell boundaries were drawn by hand on a Phalloidin/Hoechst composite image in FIJI and saved as regions of interest (ROI). Phalloidin channel images were loaded into FIJI, converted to 16-bit grayscale,

and cell areas and dimensions for each ROI were extracted using the Measure function (reporting Area and Fit Ellipse). Actin fibers were detected and quantified using the LPX FIJI plugin as described in [54], with lineExtract parameters: giwsiter = 5, mdnmsLen = 8, pickup = above (10.0), shaveLen = 3, delLen = 5, and line properties for each ROI measured using LineFeature. Parallelness (a_normAvgRad) ranges from 0 (for randomly-oriented fibers) to 1 (all fibers parallel).

***tlnrd1* and *ccm2* knockdown in zebrafish.**

Morpholinos (MOs) to knock down *tlnrd1* and *ccm2* were designed and injected using standard protocols[87]. The *ccm2* morpholino has been validated to cause cardiovascular phenotypes at the 100 μM dose[53]. A custom morpholino for Tlnrd1 (TTCCCCGAGCCACTACTAGCCATAG) was designed to target the translation start site and ordered from Gene Tools, LLC. The control oligo is a single sequence, CCTCTTACCTCAGTTACAATTTATA, that is a validated negative control[87]. Wildtype zebrafish embryos were injected with 3 nl of diluted morpholinos at multiple concentrations (50 μM, 100 μM, 200 μM, 300 μM, of control, *tlnrd1* or *ccm2* morpholinos) at the one cell stage, using a pico-injector (Harvard Apparatus). For coinjection, *tlnrd1* and *ccm2* MOs were mixed to give 50 μM of each, and 3nl of the mixture was injected. Embryos were observed for mortality and visible phenotypes at 2 days post-fertilization (dpf) and 3 dpf using a light microscope. Images were captured at 3 dpf on an EVOS microscope (Life technology) and Zeiss Axio-observer Z1. *klf2b* expression was measured by qRT-PCR on RNA isolated from 100 μM *tlnrd1* morpholino embryos at 3 dpf, using primers F: GAAGAGACACCTGTGAGGGC & R: GGACACCGATTCGTAGGACC.

**In situ hybridization for *tlnrd1* expression in zebrafish.** In situ hybridization was performed using previously validated methods[88]. Briefly, a 437 bp fragment of *tlnrd1* was amplified from genomic DNA using the PCR primers, F: CATTAACGGAATGGCAGGCG and R: TGCCCGGATAAAGGCAAAGT, subcloned and verified by sequencing. Antisense *in situ* hybridization probes were generated using an M13 reverse primer with SpeI-linearized plasmid, while sense (negative control) probes were generated using an M13 forward primer with NotI-linearized plasmid. *In situ* hybridization of embryos was conducted at 24 and 72 hrs post-fertilization using these anti-sense or sense (control) probes against *tlrnd1*.

#### Data Availability

Raw and processed data for Perturb-seq, ATAC-seq, H3K27ac ChIP-seq, and RNA-seq in TeloHAEC were deposited in NCBI's Gene Expression Omnibus under accession number GSE210523. This superseries is composed of subseries: GSE210489 (ATAC-seq), GSE210491 (ChIP-seq), GSE210522 (bulk RNAseq), GSE212396 (pilot scRNA-seq studies) and GSE210681 (comprehensive Perturb-seq). For pre-publication access, please use reviewer token: ovmhuggwtdwftez.

#### Code Availability

Data was processed using all packages required for running cNMF[29]. We additionally used R-3.6.3[89], edgeR 3.28.0[65,69,71,72], seqLogo 1.52.0, MAST 1.12.0[70], clusterProfiler 3.14.0[73,90], org.Hs.eg.db 3.10.0, Seurat 3.0.2[91], SeuratObject 4.0.0[91–93], and SingleCellExperiment 1.8.0[94] for downstream analysis in R. The supporting packages in R use for data processing and figure generation are ggplot2 3.3.5[95], ggpubr 0.4.0, ggrepel 0.9.1, gplots 3.1.1, gridExtra 2.3, scales 1.1.1, cowplot 1.1.1, dplyr 1.0.7, tidyr 1.1.3, textshape 1.7.1, reshape2 1.4.4, stringi 1.7.5[96], conflicted 1.0.4, data.table 1.14.0, purrr 0.3.4, readxl 1.3.1, writexl 1.5.0, ramify 0.3.3, optparse

1.6.6, and all dependencies. Data was further processed using pysuspenders 0.2.6[97], pysam 0.19.1[98], python 2.7.15, and LDSC v1.0.1[99]. For gene prioritization to create Supplementary Table 3.2, R 3.6.1 was used.

#### Acknowledgements

This work was supported by the Variant-to-Function Initiative at the Broad Institute (to R.M.G. and J.M.E.); NHLBI R01HL159176 (to J.M.E. and R.M.G.); the NHGRI Impact of Genomic Variation on Function Consortium (UM1HG011972 to J.M.E.); a NHGRI Genomic Innovator Award (R35HG011324 to J.M.E.); Gordon and Betty Moore and the BASE Research Initiative at the Lucile Packard Children's Hospital at Stanford University (J.M.E.); a NIH Pathway to Independence Award (K99HG009917 and R00HG009917 to J.M.E.); the Harvard Society of Fellows (J.M.E.); the Broad Institute (E.S.L.), NIH HL70567 (D.M.), and Florida Department of Health Cancer Research Chair's Fund Grant 3J-02 (to D.M.). We thank members of the Engreitz and Gupta research groups for discussions and technical assistance, and Joyce Bischoff and Sana Nasim (Harvard Medical School), for assistance with the TEER assay. E.S.L is currently on leave from the Broad Institute, MIT, and Harvard.

#### Author contributions

G.S. developed and implemented the systematic Perturb-seq method. G.S., B.C., E.S.L., R.M.G., and J.M.E. designed Perturb-seq experiments. H.K., G.S., X.R.M., T.Z., S.K.V., A.B., H.K.F., and J.M.E. developed and implemented analysis methods for Perturb-seq data. G.S., O.S-G, V.L-K and S.K.V conducted and analyzed Cell Painting experiments. G.S., V.L-K., and R.Z. conducted additional assays in endothelial cells. S.F. performed the Co-IP experiments. G.S., D.T.B., T.H.N. conducted bulk RNA-seq, ATAC-seq and ChIP-seq experiments. G.S. and K.G. analyzed bulk RNAs-seq, ATAC-seq and ChIP-seq data. P.G. and G.M. created plasmids. N.C. and H.K.F. contributed to the PoPS analysis. K.A. provided GWAS data. R.S.A. and D.M.

performed the zebrafish experiments. R.M.G. and J.M.E. supervised the work. All authors contributed to writing the manuscript.

#### Competing interests

J.M.E. is a shareholder of Illumina, Inc. and 10X Genomics. J.M.E. has received materials from 10X Genomics unrelated to this work. All other authors declare no competing interests.

#### Additional Information

Supplementary Information is available for this paper.

#### Extended Data Note 1. V2G2P approach and design considerations

We and others have previously shown that combining both "top-down" information from gene programs and "bottom-up" approaches linking variants to genes can achieve higher specificity than either category of information alone[3,25,26]. The variant-to-gene-to-program (V2G2P) approach expands upon these previous approaches by (i) generating variant-to-gene and gene-to-program maps in the same cell type; (ii) generating gene-to-program maps using Perturb-seq; and (iii) providing an interpretable, testable hypothesis linking a specific variant to a gene to a program in a given cell type. To implement this approach, we, first, constructed genome-wide enhancer-to-gene maps in endothelial cells by applying the Activity-by-Contact (ABC) model, which we recently showed performs well at linking noncoding variants to target genes in specific cell types[7,8]. Second, we created a catalog of gene programs and their regulators in endothelial cells by applying Perturb-seq to systematically study all genes in CAD GWAS loci. Perturb-seq, which involves knocking down hundreds to thousands of genes in parallel and measuring their effects on gene expression using single-cell RNA-seq, has previously been shown to provide a high-content, unbiased view of cellular programs as represented in gene expression[9–11]. Finally, we examined whether CAD genes might converge on particular gene programs by integrating gene-to-program information from Perturb-seq with variant-to-gene linking approaches.

Our approach to building a gene-to-program map using CRISPRi-Perturb-seq involved particular design considerations:

(i) We aimed to study an endothelial model relevant to the genetics of coronary artery disease. We chose telomerase immortalized human aortic endothelial cells (TeloHAEC) for these studies, because, while immortalized, they maintain important *in vitro* EC functions such as tubing, lipid transport and response to inflammatory stimuli[59,60]. We identified enhancers in resting and stimulated teloHAECs and confirmed that these regions were enriched for heritability for CAD (**Extended Data Fig. 1a, Supplementary Table 2**).

(ii) We aimed to identify cellular programs and their related genes in an unbiased manner, such that we could look for enrichment of candidate CAD genes across a range of different endothelial cell pathways. This is in contrast to the approach of selecting a particular cellular phenotype (such as endothelial cell adhesion) that may or may not be important for the genetics of disease. Accordingly, we selected Perturb-seq due to its ability to perturb many hundreds or thousands of genes in parallel, and its ability to read out the effects on all genes in the genome, thereby providing a high-throughput and high-content readout of cell states.

(iii) We aimed to design our Perturb-seq study to include all expressed nearby genes in CAD GWAS loci, as opposed to selecting just a prioritized subset of genes. This allowed us to compare various enrichment approaches, such as testing whether any nearby gene, as opposed to just nearby genes with a V2G link, were enriched in particular programs.

(iv) We aimed to perturb genes in a way consistent with presumed effects of noncoding variants, which are thought to lead to quantitative changes in the expression of genes (rather than completely eliminating expression), and which might act over long periods of time to affect disease risk. Accordingly, we used CRISPRi to quantitatively knock down gene expression (average: 40% reduction). We then read out the effects after 5 days of doxycycline induction, to allow perturbations to propagate through the network and identify how perturbations affect stable gene expression programs.

#### Extended Data Note 2. V2G2P to link disease variants to causal genes and programs

In combining V2G and G2P maps, we made several observations that help to explain the ability of V2G2P to identify disease-associated programs and genes:

(i) The intersection of V2G and G2P maps in endothelial cells was important for the identification of disease-associated programs related to endothelial functions. For 195 of 228 non-lipid signals, gene-to-program links identified more than 1 nearby gene (and up to 25) (**Extended Data Fig. 7b**), spanning all 50 programs—consistent with the notion that, by chance, a GWAS signal will have multiple nearby genes in various housekeeping and/or EC-specific programs. Statistical tests for enrichment of V2G linked-genes in programs, however, identified only 5 CAD-associated programs, all of which were endothelial cell-specific (**Fig. 2a**).

(ii) The intersection of V2G maps with CAD-associated programs was important to identify CAD-associated V2G2P genes and nominate single causal genes associated with GWAS signals. Among the 125 signals that had at least 1 V2G link, 119 signals were linked to more than 1 gene (and up to 5)—in large part due to noncoding variants being predicted to regulate more than one gene (**Extended Data Fig. 7a**), consistent with previous observations[8,28]. By contrast, of the 43 signals associated with CAD-associated V2G2P genes (V2G-linked genes in CAD-associated programs), only 6 had more than one such gene (up to 2). For example, the intersection of V2G links and G2P links to CAD-associated programs reduced the number of likely causal genes at *20p13.1*, *10p24.33* & *17q21.3* GWAS signals, where V2G and/or G2P links, individually, predicted multiple possible genes (**Extended Data Fig. 8**). We conclude that the V2G2P approach substantially refined the list of candidate disease genes compared to using V2G or G2P approaches alone (**Fig. 2c**, **Extended Data Fig. 8**).

(iii) The cell-type specificity of V2G links appeared to be important for identifying disease-associated programs. When we used a cell-type agnostic V2G approach in the V2G2P analysis (linking risk variants to the two closest genes, coding variant-containing genes, and two genes with strongest ABC links to enhancers in *any* cell type, as opposed to just endothelial cells), we found enrichment for 3 additional ubiquitous or non-endothelial cell specific processes: Program 5 (Interferon response), 36 (Steroid hormone response), and 37 (Redox homeostasis) (**Extended Data Fig. 7c**). Similarly, when we used MAGMA, which links variants to genes based on a weighted function of distance, without regard to cell-type-specific information, we also found enrichment for additional programs corresponding to processes not specific to endothelial cells (**Extended Data Fig. 6a**).

(iv) Defining programs with Perturb-seq appeared to be important. In one baseline analysis, we applied cNMF to define programs based only on the unperturbed cells in the experiment (5,506 cells carrying negative control guides), and repeated the V2G2P analysis. We found none of the programs derived from unperturbed cells were significantly enriched, and the top program included only 10 genes with V2G links instead of 18 for the top program derived from the full Perturb-seq dataset. This suggests that the scale and/or perturbations present in the full Perturb-seq experiment were important for discovering disease-associated programs and genes (**Extended Data Fig. 7e,** see Methods). In a second analysis, we computed the V2G2P enrichment using only the co-regulated genes from the Perturb-seq programs (excluding the regulators in each program), and found only Program 8 and 39 to be significant (FDR < 0.05, **Extended Data Fig. 7f**). Furthermore, most of the regulators of these programs, including CCM2, were not identified as CAD-associated V2G2P genes, because they are not co-expressed with the program. This analysis supports that Perturb-seq was important for discovering genes and programs associated with CAD.

Altogether, our results indicate that cell-type specific variant-to-gene and gene-to-program maps can be combined to effectively prioritize disease-associated programs and genes.

#### Extended Data Note 3. Additional CAD genes linked to the CCM pathway

Examining the CAD-associated V2G2P genes downstream of *CCM2* revealed insights into unresolved GWAS loci beyond the *TLNRD1* locus, and highlighted the utility of combining variant-to-gene and gene-to-program maps.

*PREX1*: At a GWAS signal at *20p13.1*, V2G2P analysis identified 2 genes that were linked by enhancer maps to a noncoding CAD variant (rs2004772) and 2 genes that were members of CAD-associated programs. Only one gene, *PREX1*, satisfied both criteria (**Extended Data Fig. 8a**). *PREX1* encodes a Rac guanine nucleotide exchange factor known to regulate actin organization[100], similar to other CCM pathway members, and is down-regulated upon *CCM2* knockdown (**Fig. 3c**). Knockout of *PREX1* has been shown to affect endothelial cell migration *in vitro* and increase vascular barrier integrity *in vivo*[57], consistent with a potential role in atherogenesis.

*SH3PXD2A* and *SLK*: At a GWAS signal at *10p24.3*, noncoding variants located in the intron of *STN1* were predicted to regulate two different CAD-associated V2G2P genes, *SH3PXD2A* and *SLK*, which were both co-regulated genes in Program 8 (**Extended Data Fig. 8b**, **Fig. 2b**). Interestingly, *SH3PXD2A* encodes an adapter protein involved in invadopodia and podosome formation[101], and *SLK* encodes a kinase that localizes to podosomes during cell migration[102], suggesting that genetic risk variants at this locus might regulate two genes with related functions.

*GOSR2*: At *17q21.3*, the noncoding variant rs17608766 has been associated with CAD risk and also with other cardiovascular phenotypes including congenital heart defects[103] and cardiac structure[104–106]. We previously linked this variant via an endothelial cell enhancer to *GOSR2*[107], which encodes a trafficking membrane protein responsible for intra-Golgi transport. Here we observed that *CCM2* knockdown led to up-regulation of *GOSR2* (+122%, $P = 8.9 \times 10^{-7}$, **Supplementary Table 19**), and *GOSR2* knockdown led to up-regulation of Program 30 (ER stress response; +94%, FDR = $2.47 \times 10^{-47}$) and down-regulation of the CAD-associated Program 35 (Focal adhesions, JUN; –23%, FDR = $8.78 \times 10^{-4}$, **Extended Data Fig. 8c**). This identifies a transcriptional phenotype for *GOSR2* in endothelial cells and suggests that *GOSR2* expression is linked to the CCM complex and other CAD-associated V2G2P genes.

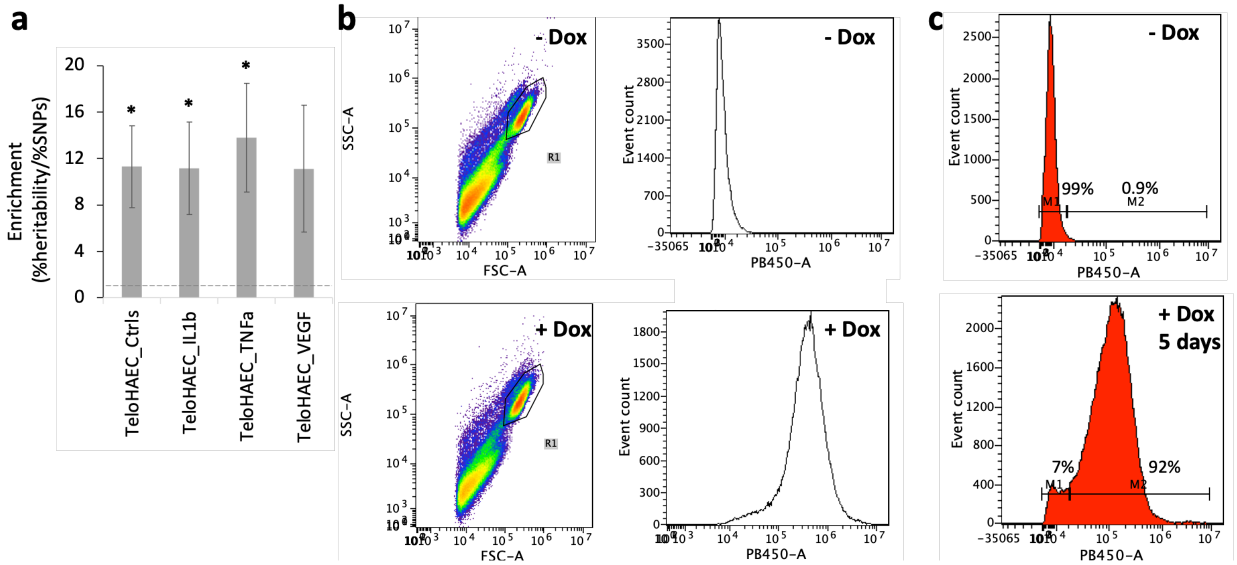


**Extended Data Fig. 1. Establishing TeloHAEC CRISPRi model**

**a)** Enrichment of CAD heritability in TeloHAEC enhancers, from Stratified Linkage Disequilibrium Score Regression analysis (S-LDSC, see Methods), where enrichment is the percentage of heritability explained by variants in enhancers (%heritability), divided by the percentage of variants in enhancers (%SNPs). Enhancers in TeloHAEC (treated under the indicated conditions) were identified from ATAC-seq and H3K27ac ChIP-seq data by the Activity-by-Contact model. Bars: standard error. *: FDR<0.05, Fisher's exact test.

**b)** FACS showing dox inducibility of KRAB-dCas9-IRES-BFP in TeloHAEC, after sorting but before the screen. Left panels: gating for viable individual cells. Right panels: Counts of gated cells by fluorescence intensity in the BFP/PB450 channel.

**c)** BFP channel counts of cells grown in parallel and concurrently with cells for the Perturb-seq screen. After expansion to 120M cells, transduction, selection and 5-day doxycycline treatment, 92% of cells remain BFP positive.

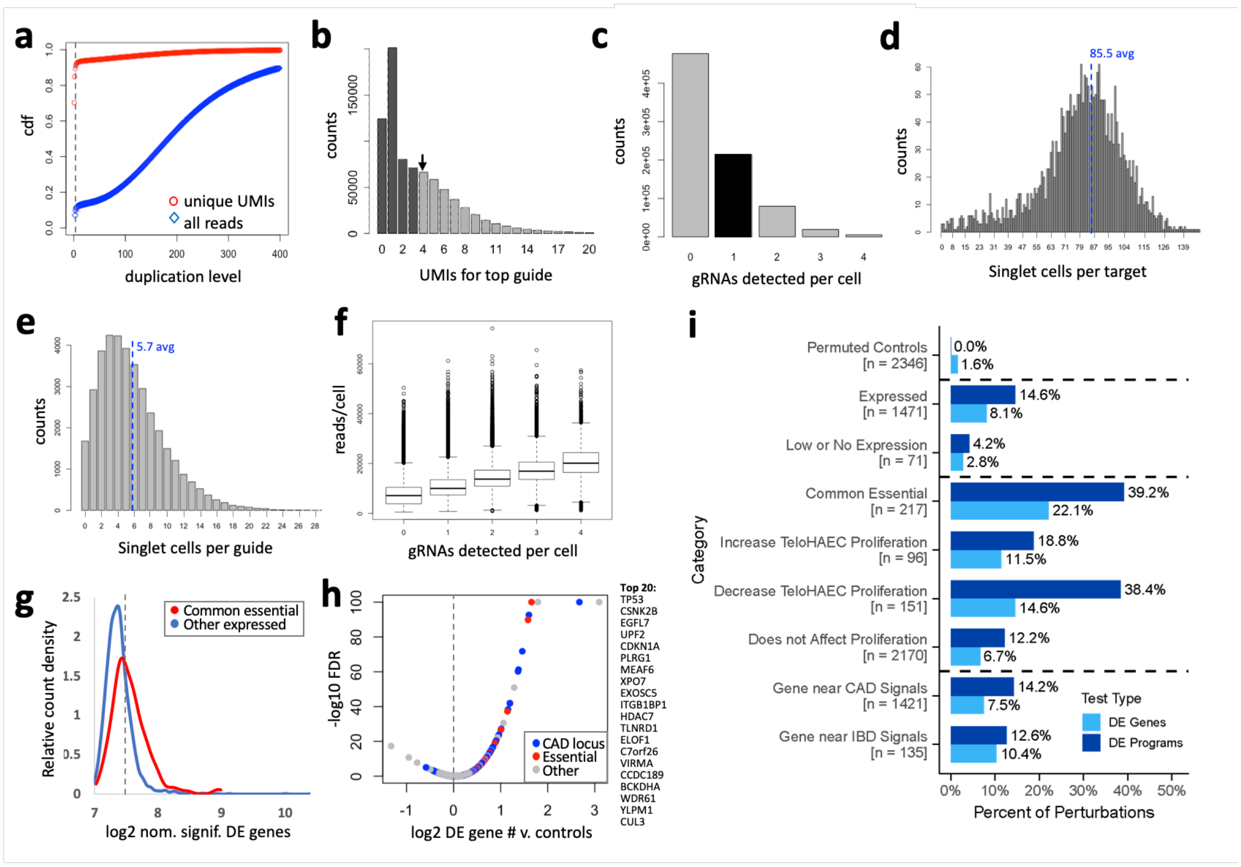


**Extended Data Fig. 2. Identification and characterization of singlets from the comprehensive CAD Perturb-seq screen.**

**a**) Cumulative distribution fraction for duplication levels of unique CBC-UMI-Guide combinations in deeply-sequenced dialout libraries ("unique UMIs", red) or all guide reads (blue) versus duplication level. Requiring 4 duplicates (dotted line) eliminates 90% of CBC-UMI-guide combinations (likely PCR chimeras), while retaining >85% of total guide reads.
**b**) UMIs for top guide per CBC. Arrow: the chosen 4 UMI threshold.
**c**) Counts of singlets (1 gRNA, black bar), doublets (2) and higher multimers, as well as cells with no guide called (0), at the chosen thresholds of 4 UMIs for the top guide and 4 or more fold fewer for the next most frequent guide.
**d**) Histogram of counts of singlet cells per target. Dotted line: average.
**e**) Histogram of counts of singlet cells per guide. Dotted line: average.
**f**) Read UMI counts for all transcripts per cell by singlet/multiplet status. Median UMIs per singlet cell was 9,997, and average was 10,870. The median for cells with no guide called was 7,125, indicating that low guide UMI count is associated with low overall UMI count. Median UMIs for doublets was 13,723, 37.3% more than singlets. Assuming that droplets with two cells will have double the number of reads, this suggests 37% of doublets are due to two cells (9.3% of cells with guides) while the remainder (15.7% of cells with guides) are due to two guides in one cell, very close to the expectation from the infection MOI of 15%.
**g**) Number of nominally significant differentially expressed (DE) genes per perturbed target (genes with raw p <0.01, and fold change >1.15 from EdgeR DE analysis). Perturbations that affected the transcriptome were those that significantly increased the number of nominally significant DE genes relative to the 48 targeted negative control genes (not expressed in TeloHAEC). Dotted line: 95th percentile number of DE genes for negative controls. 245 perturbations had a significant effect on the transcriptome, FDR <0.05 (10.7% of all targets that were not negative controls: including 31.9% of common essential genes (red, defined as in Fig. 1d) and 9.0% of other genes (blue)).

**h**) Volcano plot showing $\log_2$ (# DE genes for target)/(avg. # DE genes for non-expressed controls) versus $-\log_{10}$ FDR (capped at 100). Right: Symbols for target genes with the strongest effects.
**i**) Percent of perturbations that have a significant transcriptional effect in Perturb-seq, as defined by either (i) “DE Genes”: perturbations with significant effect on the transcriptome, as compared to 48 non-expressed negative control promoters, by binomial test (see Methods) or (ii) “DE Programs”: perturbations that lead to significant changes in program expression by MAST with 10X lane correction (FDR < 0.05).
Permuted Controls: Simulated negative controls, where statistical tests were performed on randomly drawn cells that carry negative control or safe-targeting guides.
Expressed: Genes with >1 transcripts per million (TPM) in teloHAEC control bulk RNA-seq.
Low or No Expression: Genes with less than or equal to 1 TPM.
Common Essential: Common essential genes from DepMap[108].
TeloHAEC Proliferation: Fitness effects observed in the Perturb-seq experiment, by comparing guide frequencies (see Methods). Increase: >15% increase in guide frequency (FDR < 0.05), Decrease: >15% decrease in guide frequency (FDR < 0.05).
Gene near CAD GWAS signals: Expressed genes nearby any CAD GWAS signal (2 closest on each side, and all within +/-500kb).
Gene near IBD signals: Expressed genes nearby 10 selected IBD GWAS signals (closest 2 genes & all within +/-500kb), with no genes overlapping those for CAD signals.

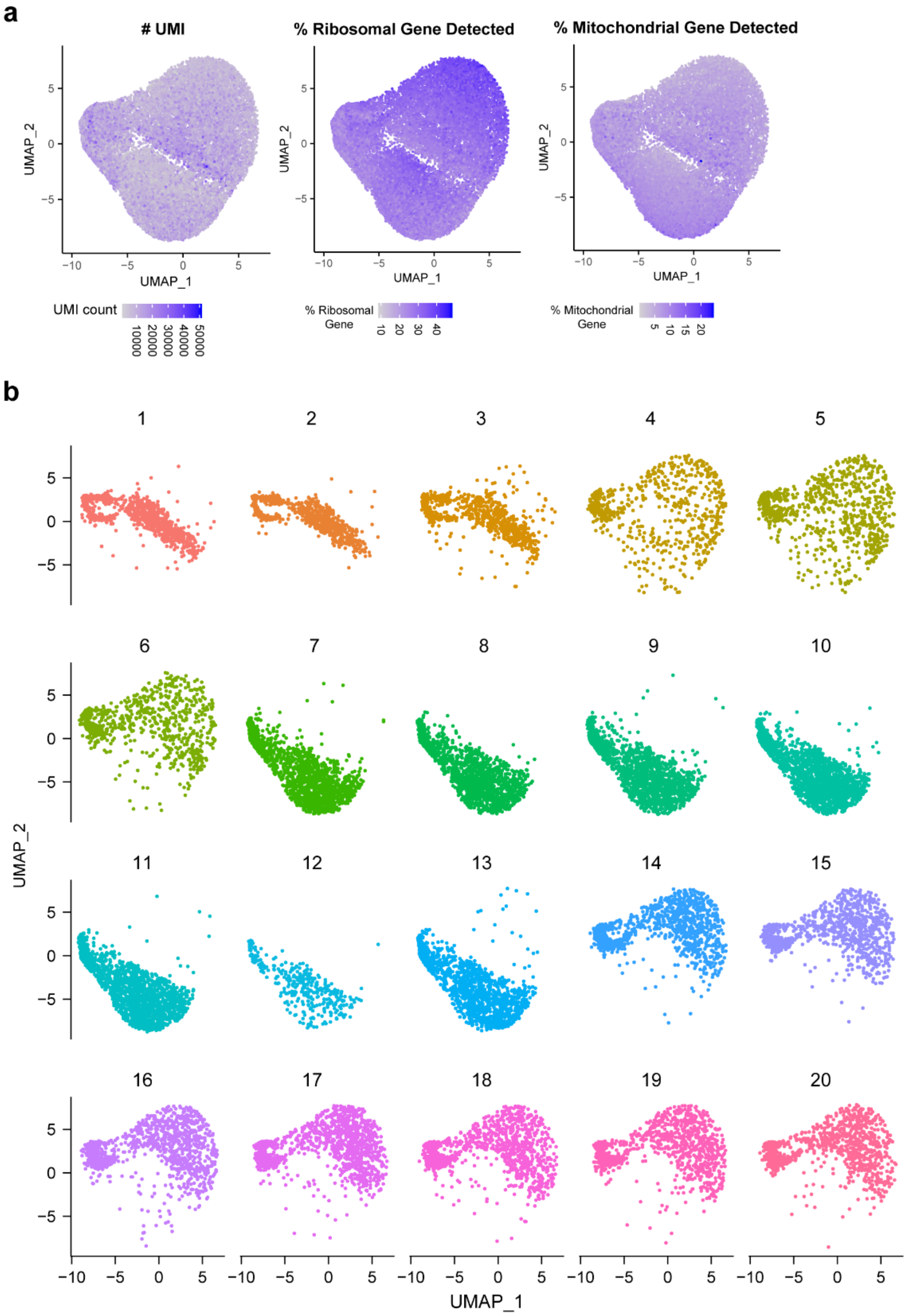


**Extended Data Fig. 3. QC metrics of all single cells in UMAPs**

**a**. UMAPs showing number of UMIs per cell (left), percent ribosomal genes detected per cell (middle), percent mitochondrial genes detected per cell (right).

**b**. UMAPs showing cells from each of the twenty 10X lanes. The differences in clustering along the UMAP_2 axis indicates a technical batch effect between 10X lanes.

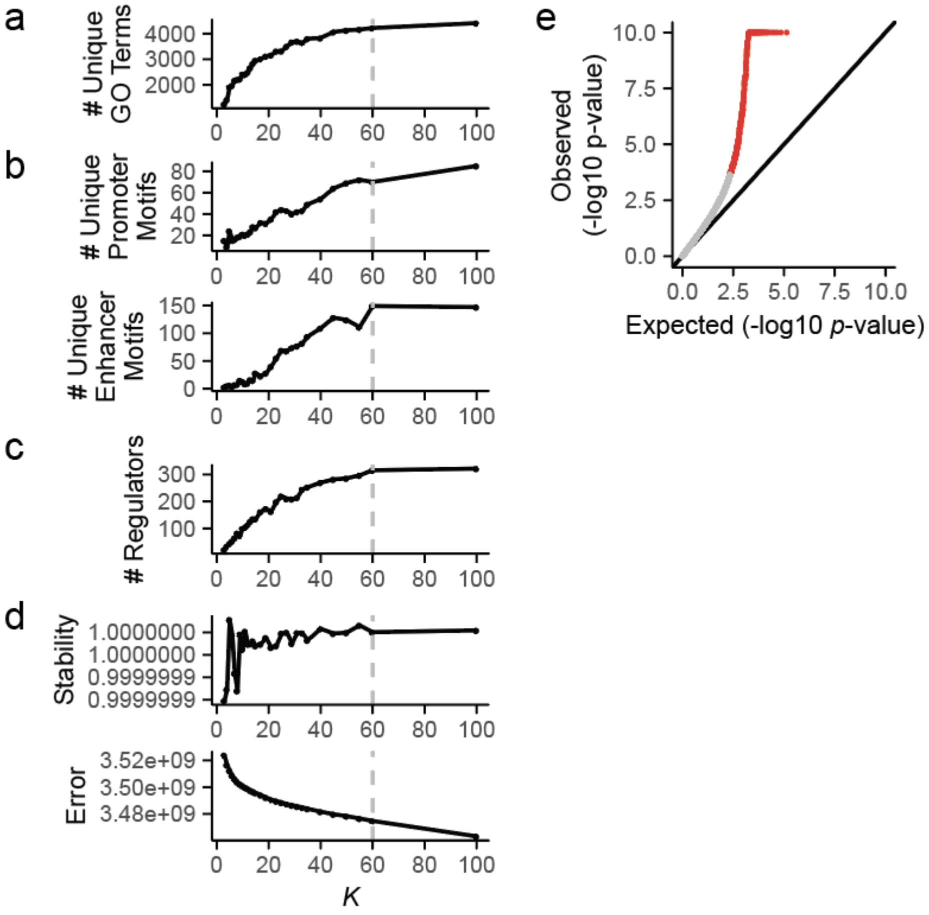


**Extended Data Fig. 4. Selecting number of components for cNMF**

**a**. Gene set enrichment analysis for GO terms among co-regulated genes, as a function of the number of components in the cNMF model (*K*). *y*-axis: The number of unique GO terms enriched across all programs for a given *K*.

**b**. Number of unique motifs enriched among the promoters (top) or enhancers (bottom) of co-regulated genes across all components, as a function of *K*.

**c**. Number of unique perturbations that have significant effect (FDR < 0.05) on one or more programs, as a function of *K*.

**d**. Model-based evaluation of the choice of *K*. Stability of the components over 100 NMF runs (top) and element-wise square of error (bottom, see Methods).

**e**. Quantile-quantile plot for effects of perturbations on program expression. X-axis: Expected uniform distribution. Y-axis: $-\log_{10}$ *p*-value from MAST. Red: *p*-value < 0.05.

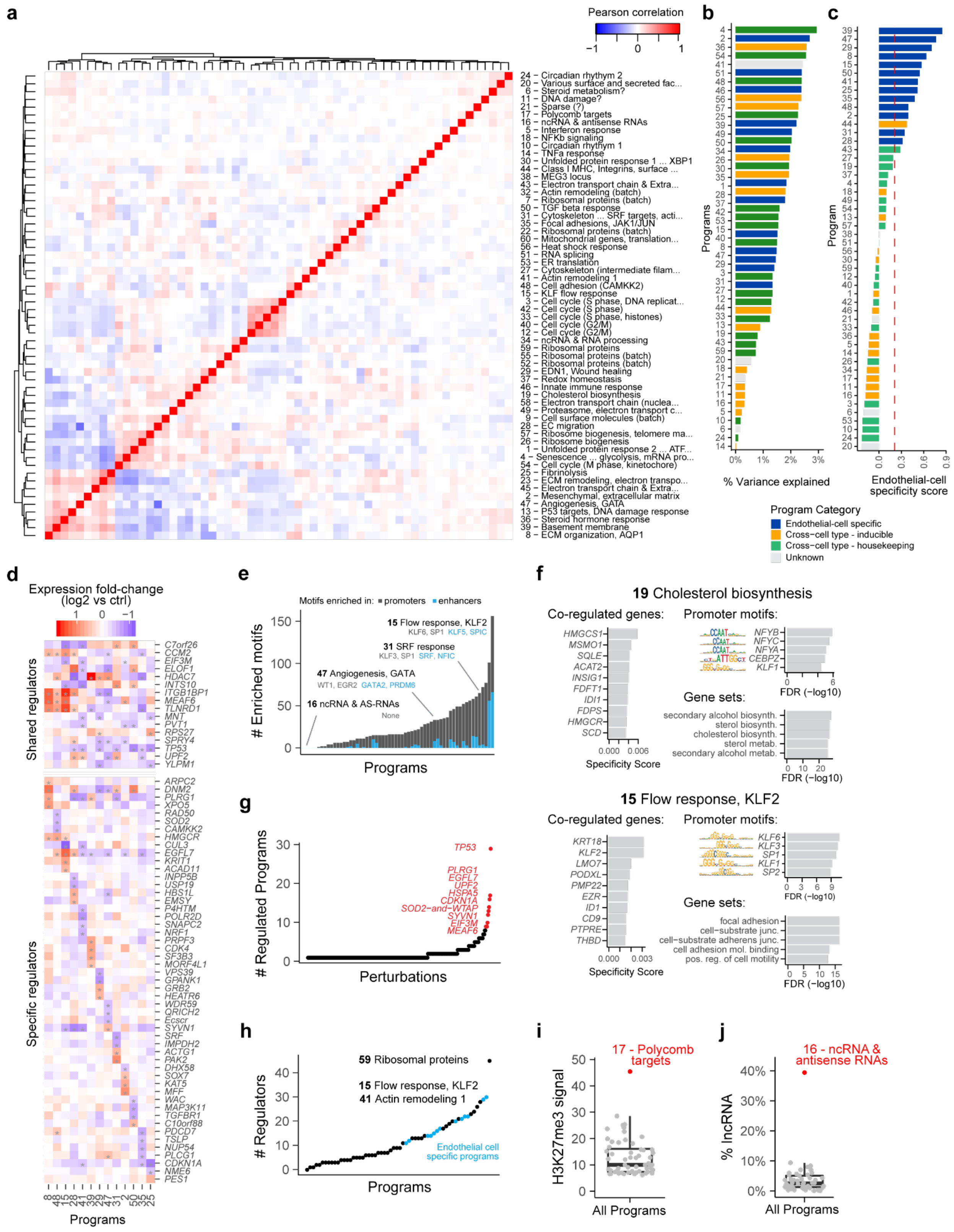

**Extended Data Fig. 5. Catalog of gene programs**

**a.** Correlation heatmap of cNMF components. Color: Pearson's correlation of $log_2$ fold-change in component expression across all perturbed genes.
**b.** 50 programs ordered by variance explained (see Methods).
**c.** 50 programs ordered by endothelial-cell specificity score — that is, the degree to which the co-regulated genes in the program are specifically expressed in endothelial cells versus in other cell types from FANTOM5 CAGE data (see Methods). Red line: *z*-score corresponding to top 10% of genes most specifically expressed in endothelial cells.
**d.** Effects of selected regulators on the 13 endothelial-cell-specific programs. Heatmap: $log_2$ fold-change in component expression in perturbation vs control. Top: 16 regulators shared between multiple endothelial cell-specific programs. Bottom: the 4 significant regulators (experiment-wide FDR < 0.05) per program with the most specific effects on that program relative to other endothelial-cell-specific programs.
**e**. Programs ordered by number of enriched transcription factor motifs (See Methods). Gray: promoters. Blue: enhancers. Some programs only have enrichment for motifs in promoters. Some programs showed enrichment of distinct motifs in enhancers versus promoters, such as Program 47 (Angiogenesis, GATA2), with promoter enrichment in WT1 and EGR2 motifs, and enhancer enrichment in GATA2 and PRDM6 motifs. Among the programs with few or no enriched transcription factor motifs, we identified other likely proximal regulatory mechanisms: Program 17 expressed genes whose promoters were marked by H3K27me3 in endothelial cells (see also **Extended Data Fig. 5i**), and the most significant regulator of this program was *SUZ12*, a component of the complex (PRC2) that writes this histone modification; and Program 16 pointed to a potential RNA surveillance program, since 40% of its program genes were noncoding RNAs (**Extended Data Fig. 5j**), and its regulators included a component of the RNA exosome (*EXOSC5*) and the chromatin remodeler *INO80E*, which has previously been shown to regulate a subset of noncoding transcripts in yeast[109] (see also **Supplementary Table 11**).
**f.** Annotations for two example programs: 19 (Cholesterol biosynthesis) and 15 (Flow response, KLF). Left: Top 10 program co-regulated genes. Right, top: Motifs enriched in promoters of the 300 program co-regulated genes. Right, bottom: Gene Ontology terms enriched in the 300 program co-regulated genes.
**g**. Perturbations ordered by the number of regulated programs. Red: top 10 genes.
**h**. Programs ordered by the number of regulators. Blue: endothelial-cell-specific programs.
**i**. Average H3K27me3 ChIP-seq signal in co-regulated gene promoters. The top program is Program 17 (Polycomb targets). See legend to (**e**) for more details.
**j.** Percent of noncoding RNA genes in program co-regulated genes. The top program is Program 16 (ncRNA & antisense RNAs). See legend to (**e**) for more details.

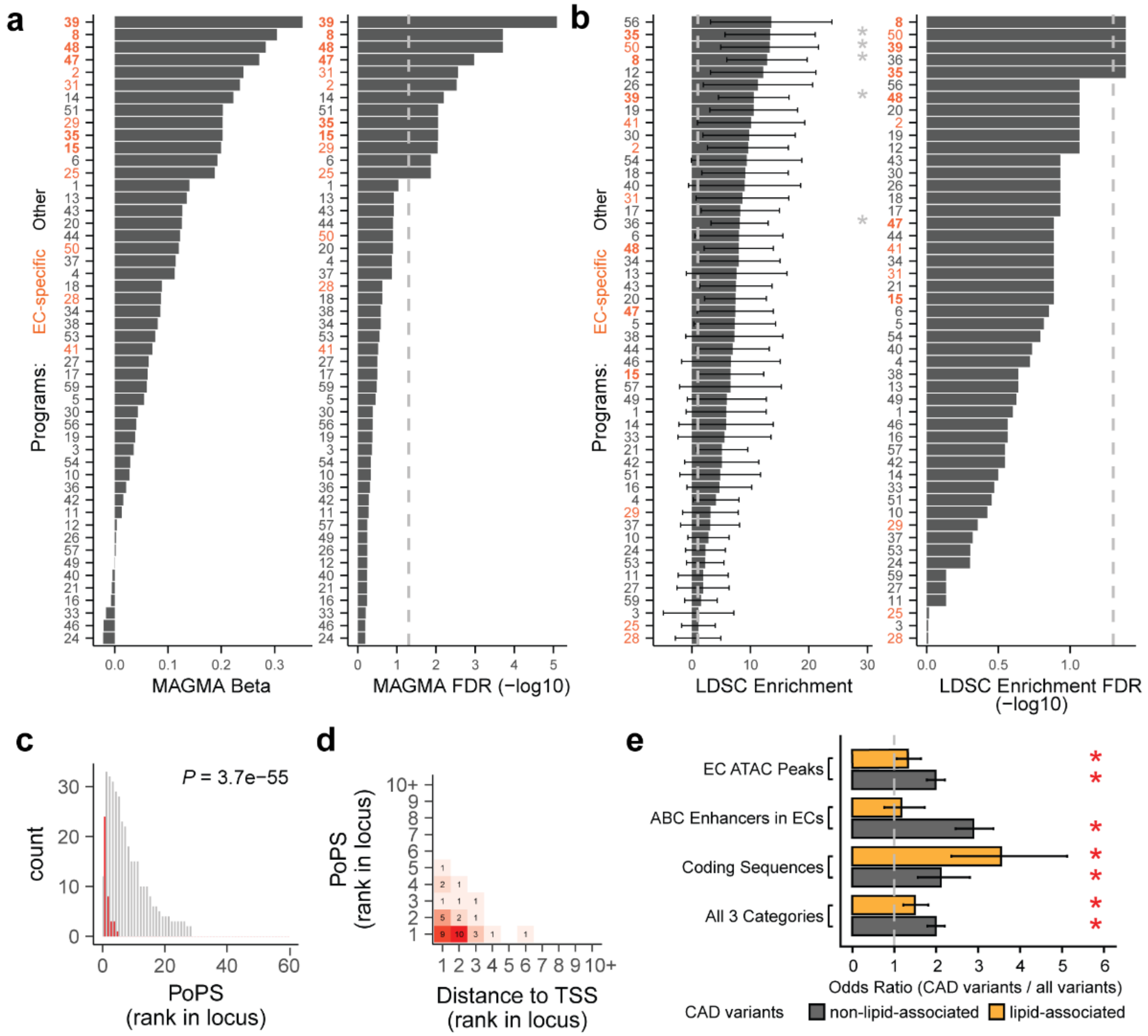


**Extended Data Fig. 6. Prioritization of CAD-associated programs and candidate CAD genes**

**a.** Using MAGMA to prioritize gene programs enriched for CAD heritability (linking variants to program genes and 50kb of flanking sequence, see Methods). Barplots show beta regression coefficient (left) and $-\log_{10}$ FDR (Benjamini-Hochberg adjusted enrichment *p*-value, right). Programs are ordered separately by beta or FDR value. Dotted line: FDR = 0.05.

**b.** Using S-LDSC to prioritize gene programs enriched for CAD heritability (linking variants in endothelial cell chromatin accessible regions to genes within 50 Kb, see Methods). Barplots show enrichment (left) and $-\log_{10}$ FDR (Benjamini-Hochberg adjusted enrichment *p*-value, right). Error bars: 95% confidence interval for the mean. *: FDR < 0.05. Dotted line: 1 fold enrichment (left), or FDR 0.05 (right).

**c**. CAD-associated V2G2P genes are ranked highly by an independent gene prioritization method, the Polygenic priority score (PoPS). For each of the 43 CAD GWAS signals including a CAD-associated V2G2P gene, we ranked nearby genes based on their PoPS scores. Red: 39 CAD-associated V2G2P genes (2 genes, *EXOC3L2* and *PECAM1*, were not assigned scores by PoPS). Gray: all other nearby genes. *p*-value: Mann-Whitney U-test.

**d**. Contingency table of PoPS and distance-to-TSS ranks for the 39 CAD-associated V2G2P genes. (2 CAD-associated V2G2P genes were not assigned scores by PoPS).

**e.** Odds ratios of variants in lipid-associated or non-lipid-associated CAD GWAS signals in (i) ATAC peaks in endothelial cells, (ii) ABC enhancers in endothelial cells, (iii) coding sequences, or (iv) all three categories combined, compared to background variants (all SNPs from 1000 Genomes, see Methods). Bar plots: odds ratios (CAD vs control variants). Error bars: 95% confidence interval. *: p-value < 0.05, Fisher's exact test.

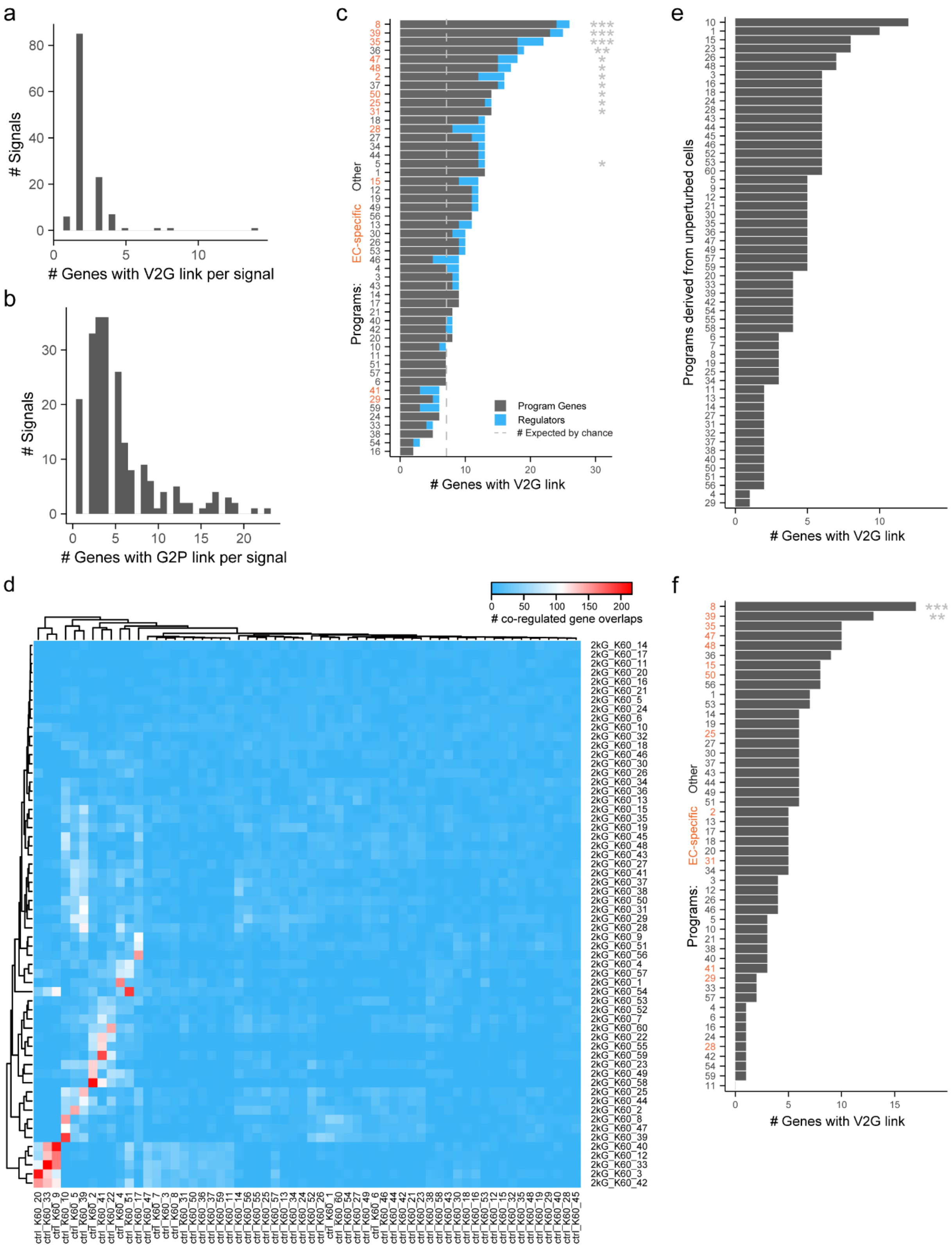


**Extended Data Fig. 7. Details for V2G2P analysis**

**a.** Number of genes with V2G links, per non-lipid CAD GWAS signal.

**b.** Number of genes with G2P links, per non-lipid CAD GWAS signal.
**c.** The cell-type specificity of V2G links appeared to be important for identifying endothelial-cell-specific programs. Here, we repeated the V2G2P analysis (as in **Fig 2a**), but linked variants to genes using cell-type-agnostic criteria (including ABC scores from any cell type and not just endothelial cells). The 50 programs are ordered (*y*-axis) by the number of program genes linked to CAD variants (*x*-axis). Gray dashed line: the number of genes linked to CAD variants that would be expected by chance. Orange labels: endothelial-cell-specific programs. Two significant non-endothelial-cell-specific programs were identified. (**FDR* <0.05, ***FDR* <0.005, ****FDR* <5e-4)
**d**. Number of overlapping co-regulated genes between control programs (ctrl, x-axis) and full library (2kG, y-axis) programs.
**e**. Using the full Perturb-seq dataset appeared to be important for identifying the 5 CAD-associated programs. Programs discovered through cNMF analysis of only the “unperturbed” cells carrying negative control guideRNAs. In this version of the analysis, none of the programs are enriched for genes with V2G link.
**f**. Enrichment of genes with V2G links, from the full library but only using co-regulated genes (not regulators,***: FDR < 0.0005, **: FDR < 0.005)

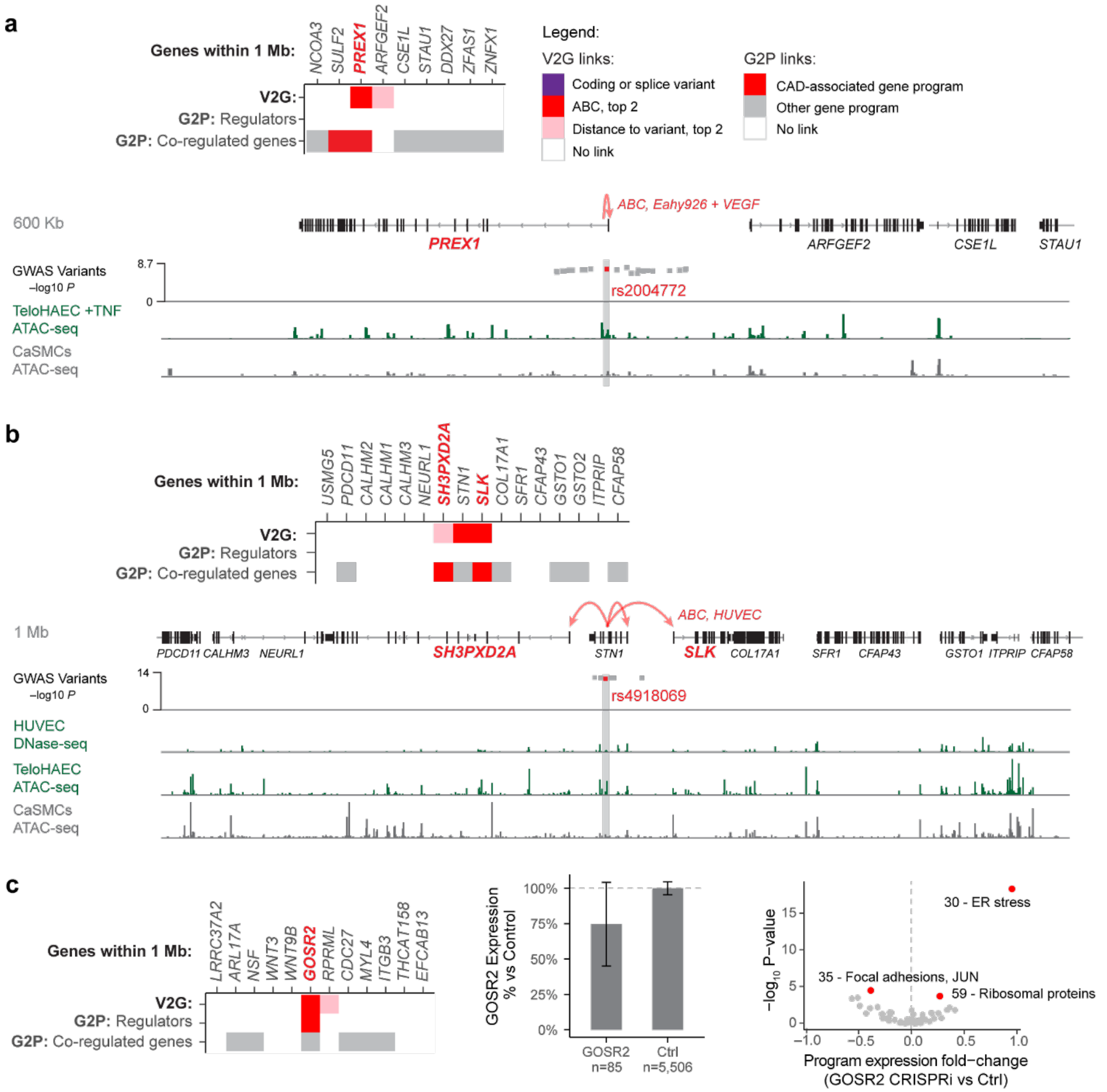


**Extended Data Fig. 8. Variant-to-gene to program links refine causal gene predictions.**
**a.** V2G2P evidence at the *20p13.1* CAD GWAS locus. *Top:* Heatmap lists genes within 1 Mb of the CAD GWAS signal in genomic order, and shows variant-to-gene (V2G) and gene-to-pathway (G2P) evidence, with the prioritized CAD-associated V2G2P gene(s) labeled in red bold font. Legend details: "ABC, top 2": A noncoding variant overlaps a chromatin accessible peak in endothelial cells, and the ABC score is at least the second highest of all genes near the GWAS signal. "Distance to variant, top 2": A noncoding variant overlaps a chromatin accessible peak in endothelial cells, and the gene is one of the two closest genes to the variant. *Bottom:* Zoom-in on genes near the CAD GWAS signal, where rs2004772 is predicted by ABC to regulate *PREX1* in the Eahy926 endothelial cell line treated with VEGF. Red dot: Prioritized variant in predicted enhancer. Gray dots: Other variants within $R^2$ < 0.9 of the lead variant in the locus. Signal tracks below show ATAC-seq or DNase-seq for endothelial and coronary artery smooth muscle cells (CaSMCs, another cell type relevant to CAD).
**b.** As per **a**, showing V2G2P evidence at the *10p24.33* CAD GWAS signal, where three genes had V2G links (to an enhancer containing rs4918069) and 2 had gene to CAD-associated program links. HUVEC: human umbilical vein endothelial cells.

**c.** V2G2P evidence at the *17q21.3* CAD GWAS locus, where we have previously linked rs17608766 to *GOSR2*[107]. Heatmap, as in panel (a). Middle: Barplot shows efficiency of knockdown of *GOSR2* in Perturb-seq. Error bars: 95% confidence interval. Right: Volcano plot shows effect of *GOSR2* knockdown in Perturb-seq on the expression of the 50 non-batch gene programs. Red: FDR < 0.05.

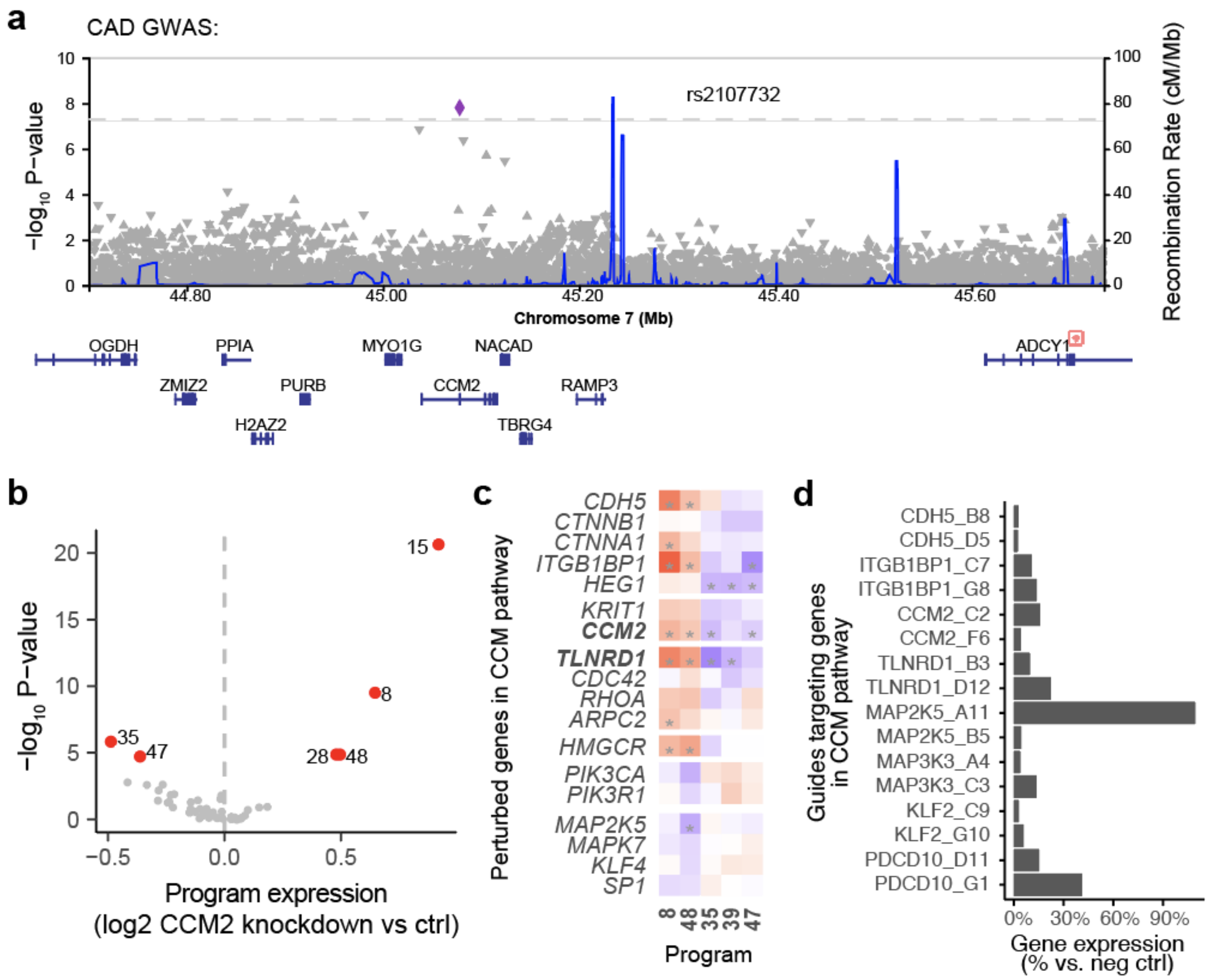


**Extended Data Fig. 9. Regulatory connections amongst perturbed genes and the CCM pathway**

**a.** Locus zoom plot for CAD GWAS in a 1-Mb region around *CCM2*.

**b.** Volcano plot showing effect of *CCM2* knockdown in Perturb-seq on the expression of the 50 programs. Red: FDR < 0.05.

**c.** Effects of selected perturbed genes on CAD-associated programs (Same as **Fig. 3b**, with significant effects marked with a * (FDR < 0.05)). Color scale: $\log_2$ fold-change on program expression in Perturb-seq. Bold text: CAD-associated V2G2P genes.

**d.** Knockdown efficiency for genes in the CCM pathway, in bulk RNA-seq (**Fig. 3c**). x-axis: Gene expression for each target gene in cells receiving target guides, vs. cells with control guides. y-axis: guide IDs for CCM pathway genes.

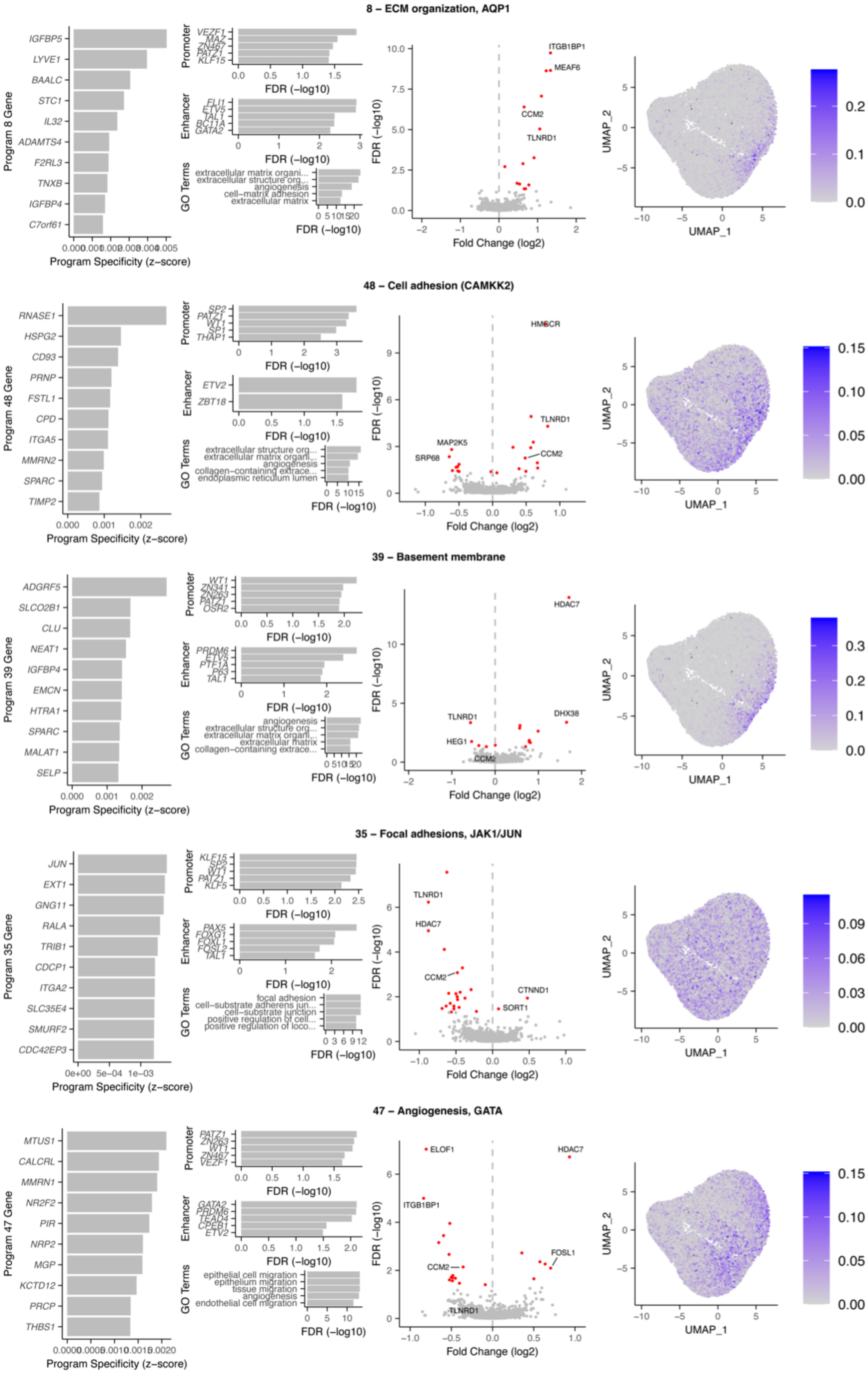

8 – ECM organization, AQP1
IGFBP5
LYVE1
BAALC
STC1
IL32
ADAMTS4
F2RL3
TNXB
IGFBP4
C7orf61
Program 8 Gene
Program Specificity (z–score)
Promoter
VEZF1
MAZ
ZN467
PATZ1
KLF15
Enhancer
FLI1
ETV5
TAL1
BC11A
GATA2
GO Terms
extracellular matrix organi...
extracellular structure org...
angiogenesis
cell–matrix adhesion
extracellular matrix
FDR (–log10)
ITGB1BP1
MEAF6
CCM2
TLNRD1
Fold Change (log2)
UMAP_1
UMAP_2
48 – Cell adhesion (CAMKK2)
RNASE1
HSPG2
CD93
PRNP
FSTL1
CPD
ITGA5
MMRN2
SPARC
TIMP2
Program 48 Gene
SP2
PATZ1
WT1
SP1
THAP1
ETV2
ZBT18
extracellular structure org...
extracellular matrix organi...
angiogenesis
collagen–containing extrace...
endoplasmic reticulum lumen
HMGCR
TLNRD1
MAP2K5
SRP68
CCM2
39 – Basement membrane
ADGRF5
SLCO2B1
CLU
NEAT1
IGFBP4
EMCN
HTRA1
SPARC
MALAT1
SELP
Program 39 Gene
WT1
ZN341
ZN263
PATZ1
OSR2
PRDM6
ETV5
PTF1A
P63
TAL1
angiogenesis
extracellular structure org...
extracellular matrix organi...
extracellular matrix
collagen–containing extrace...
HDAC7
TLNRD1
DHX38
HEG1
CCM2
35 – Focal adhesions, JAK1/JUN
JUN
EXT1
GNG11
RALA
TRIB1
CDCP1
ITGA2
SLC35E4
SMURF2
CDC42EP3
Program 35 Gene
KLF15
SP2
WT1
PATZ1
KLF5
PAX5
FOXG1
FOXL1
FOSL2
TAL1
focal adhesion
cell–substrate adherens jun...
cell–substrate junction
positive regulation of cell...
positive regulation of loco...
TLNRD1
HDAC7
CCM2
CTNND1
SORT1
47 – Angiogenesis, GATA
MTUS1
CALCRL
MMRN1
NR2F2
PIR
NRP2
MGP
KCTD12
PRCP
THBS1
Program 47 Gene
PATZ1
ZN263
WT1
ZN467
VEZF1
GATA2
PRDM6
TEAD4
CPEB1
ETV2
epithelial cell migration
epithelium migration
tissue migration
angiogenesis
endothelial cell migration
ELOF1
HDAC7
ITGB1BP1
FOSL1
CCM2
TLNRD1

#### Extended Data Fig. 10. Annotations for CAD-associated programs: 8, 35, 39, 47, 48

**Left panels.** Top 10 program co-regulated genes. Program Specificity z-scores are the cNMF marker gene coefficients, indicating how specific this gene is to this program, relative to other programs (see Methods).
**Middle left panels.** Top: Top 5 motifs enriched in the promoters or enhancers of the program co-regulated genes. Bottom: Top 5 GO terms enriched in program co-regulated genes.
**Middle right panels.** Regulators of the program. Volcano plot shows effects of all perturbed genes on program expression. Red: FDR < 0.05. Labeled: top 2 significant regulators in each direction, plus *CCM2* and *TLNRD1*.
**Right panels.** UMAP of program expression in a subset of cells (24,000, randomly selected).

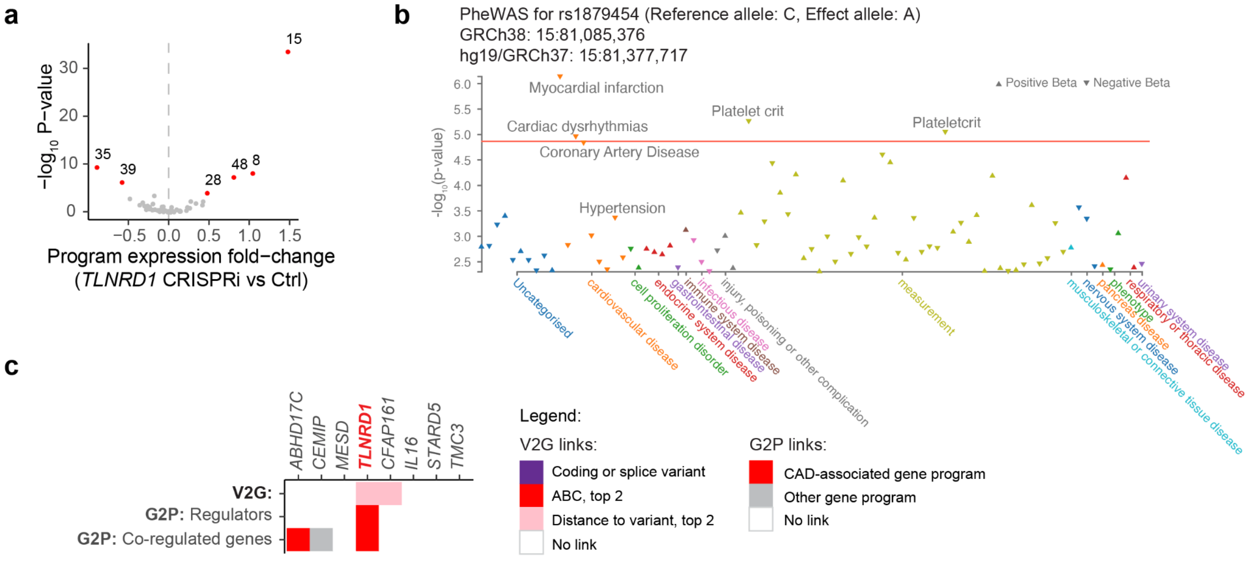


#### Extended Data Fig. 11. *TLNRD1*

**a.** Volcano plot showing the effect of *TLNRD1* knockdown in Perturb-seq on the expression of the 50 programs. Red: FDR < 0.05.

**b.** Phenome-wide association study (PheWAS) for rs1879454, from Open Targets[110,111]. No lipid measure met the p.value threshold for inclusion in the plot, of 0.005. Note: The p.value for CAD is higher than that observed in Aragam et al.[13] because Open Targets does not currently contain summary statistics for the latest CAD GWAS. There were no measures of circulating lipids or blood pressure associated with this GWAS signal in a PheWAS analysis in Aragam et al.[13].

**c.** Variant-to-gene-to-program evidence at the *15q25.1* CAD GWAS locus. Heatmap lists genes within 1 Mb of the CAD GWAS signal in genomic order. Heatmap shows variant-to-gene (V2G) and gene-to-pathway (G2P) evidence, with the CAD-associated V2G2P gene labeled in red bold font. Legend details: "ABC, top 2": A noncoding variant overlaps a predicted enhancer linked to this gene in endothelial cells, and ABC score is at least the second highest of all genes in the locus. "Distance to variant, top 2": A noncoding variant overlaps a chromatin accessible peak near this gene in endothelial cells, and the gene is one of the two closest genes to the peak. CAD-associated programs: 8, 35, 39, 47, 48.

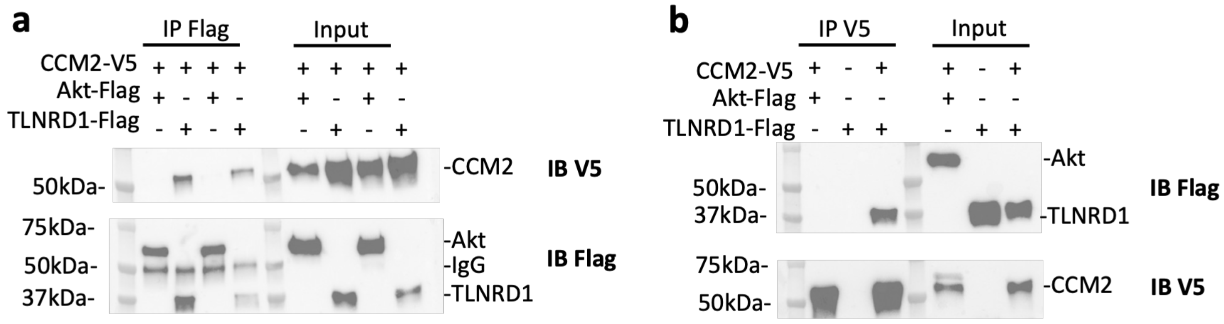


**Extended Data Fig. 12. Reciprocal co-immunoprecipitation of CCM2 and TLNRD1**

**a**. HEK293 cells were transfected with V5-tagged CCM2 and either Flag-tagged TLNRD1 or Flag-tagged Akt (negative control). Cell lysates were either immunoprecipitated with anti-Flag beads (IP Flag), or loaded directly on the gel (Input). The membranes were first probed for V5 to detect CCM2 in the Flag precipitant and confirm the transfection of CCM2-V5. The membranes were then re-blotted for Flag to evaluate the efficiency of Flag immunoprecipitation and validate the transfection of Akt-Flag and TLNRD1-Flag. Each pair of lanes came from independent biological replicates.

**b**. HEK293 cells were transfected with V5-tagged CCM2 and Flag-tagged TLNRD1, or, as negative controls, either with CCM2-V5 and Akt-Flag or only TLNRD1-Flag. Cell lysates were either immunoprecipitated with anti-V5 beads (IP V5), or loaded directly on the gel (Input). The membranes were first blotted for Flag to detect TLNRD1 in the V5 precipitant and validate the transfection of Akt-Flag and TLNRD1-Flag. The membranes were re-blotted for V5 to evaluate the efficiency of V5 immunoprecipitation and confirm the transfection of CCM2-V5.

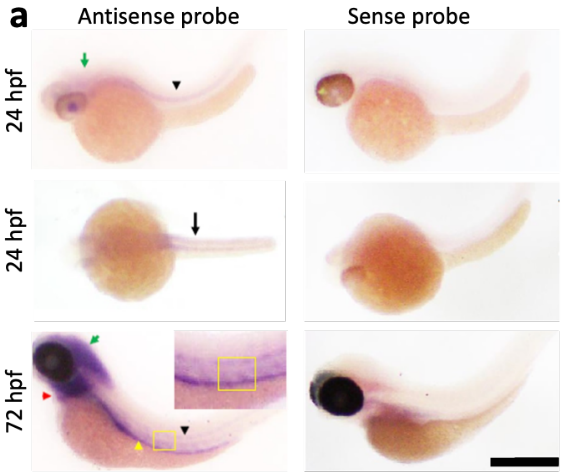


**b**

| Morpholino | Experiment | Concentration (µM) | Volume (nl) | Injected (#) | Survival @ 3dpf: #(%) | Heart Phenotype: #(%) |
|---|---|---|---|---|---|---|
| Ccm2 MO | 1 | 50 | 3 | 50 | 41 (82%) | 0 |
| Ccm2 MO | 1 | 100 | 3 | 150 | 98 (63%) | 69 (70%) |
| Ccm2+Tlnrd1 MO | 1 | 50 each | 3 | 50 | 39 (78%) | 26 (67%) |
| Ccm2+Tlnrd1 MO | 2 | 50 each | 3 | 125 | 103(85%) | 77(74%) |
| Control MO | 1 | 50 | 3 | 50 | 45 (90%) | 0 |
| Control MO | 1 | 100 | 3 | 150 | 127 (84%) | 0 |
| Control MO | 2 | 100 | 3 | 50 | 42(84%) | 0 |
| Control MO | 1 | 200 | 3 | 50 | 40 (80%) | 0 |
| Control MO | 1 | 300 | 3 | 50 | 34 (68%) | 0 |
| Non injected | 1 | NA | NA | 50 | 49 (96%) | 0 |
| Tlnrd1 MO | 1 | 50 | 3 | 50 | 42 (84%) | 0 |
| Tlnrd1 MO | 1 | 100 | 3 | 150 | 118 (78%) | 97(83%) |
| Tlnrd1 MO | 1 | 200 | 3 | 50 | 27 (54%) | 24(88%) |
| Tlnrd1 MO | 1 | 300 | 3 | 50 | 20 (40%) | Deformed |
| Tlnrd1 MO | 1 | 400 | 3 | 50 | 17 (34%) | Deformed |
| Tlnrd1 MO | 1 | 1000 | 3 | 50 | 3 (6%) | Deformed |

**Extended Data Fig. 13. Zebrafish *tlnrd1* staining & morpholino statistics**

**a.** *In situ* hybridization of embryos at 24 and 72 hours post-fertilization (hpf) using an antisense probe against *tlnrd1* mRNA (with the corresponding sense probe used as a negative control). *tlnrd1* was expressed at all time points in the head, notochords, and tail vasculature. Green arrowhead: *tlnrd1* staining in the brain, black arrowhead: notochords, red arrowhead: heart, yellow arrowhead: gut. Black arrow in the dorsal view at 24 hpf indicates the myotomes. Inset panel: enlargement of the region indicated by the yellow box. Yellow box in inset panel highlights dorsal aorta & tail vein. Scale bar = 1 mm.

**b.** Summary of morpholino studies. “MO”: morpholino. “Deformed” indicates embryos that were too misshapen to accurately assess heart phenotypes.

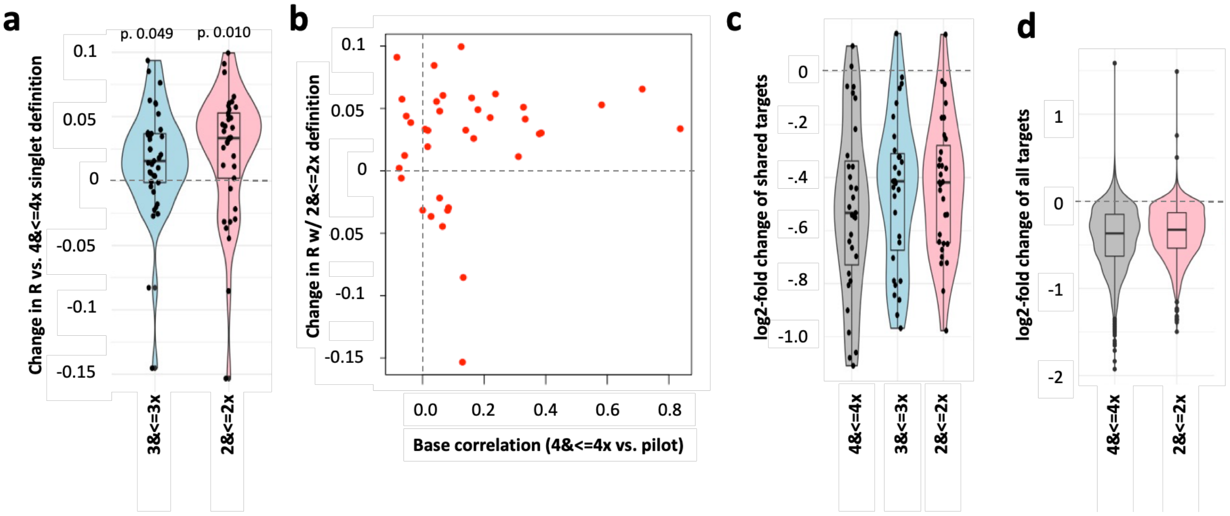


**Extended Data Fig. 14. Power and accuracy considerations for singlet thresholds**

**a**. We measured the correlation between differential expression log2 fold changes resulting from 37 perturbed gene targets shared between a pilot library and our full-scale library. Violin plots show the increases in correlation coefficients (*R*) between relaxed threshold comparisons (pilot v. full library 3<=3x, and pilot v. full library 2<=2x) and the base comparison (pilot vs. full library 4<=4x), where singlet thresholds are abbreviated as "[UMIs for the top guide required to assign a guide to a cell] <= [fold lower number of UMIs for the 2nd to top guide, to assign a singlet]".

**b**. Plot of the change in *R* for each target using the full library 2<=2x singlet definition vs. the 4<=4x singlet definition (Y axis, same as rightmost violin plot in (a)) against the *R* value for the base correlation (between the pilot and the 4<=4x full library singlet definition, X axis).

**c**. Violin plots of log2 fold changes for knock down of the target genes in (a), in the full-scale library, for each singlet definition. Medians: -0.53 for 4<=4x, -0.41 for 3<=3x and -0.42 for 2<=2x.

**d**. As in (c), but for all 2885 targets of the full-scale library. Median log2fc for targets with the 4<=4x singlet definition was -0.368, and with the 2<=2x singlet definition was -0.327.
